# CONCERTED PLANT GROWTH AND DEFENSE THROUGH TARGETED PHYTOHORMONE CROSSTALK MODIFICATION

**DOI:** 10.1101/2024.04.08.588615

**Authors:** Grace A. Johnston, Hannah M. Berry, Mikiko Kojima, Hitoshi Sakakibara, Cristiana T. Argueso

**Affiliations:** Department of Agricultural Biology, Colorado State University, Fort Collins, CO 80523, USA; Graduate Program for Cell and Molecular Biology, Colorado State University, Fort Collins, CO 80523, USA; RIKEN Center for Sustainable Resource Science, Suehiro, Tsurumi, Yokohama 230-0045, Japan; Graduate School of Bioagricultural Sciences, Nagoya University, Chikusa, Nagoya, 464-8601 Japan

**Keywords:** Cytokinin, salicylic acid, jasmonic acid, phytohormone crosstalk, growth-defense tradeoff

## Abstract

Plant immunity activation often results in suppression of plant growth, particularly in the case of constitutive immune activation. We discovered that signaling of the phytohormone cytokinin (CK), known to regulate plant growth through the control of cell division and shoot apical meristem (SAM) activity, can be suppressed by negative crosstalk with the defense phytohormones jasmonic acid (JA), and most evidently, salicylic acid (SA). We show that changing the negative crosstalk of SA on CK signaling in autoimmunity mutants by targeted increase of endogenous CK levels results in plants resistant to pathogens from diverse lifestyles, and relieves suppression of reproductive growth. Moreover, such changes in crosstalk result in a novel reproductive growth phenotype, suggesting a role for defense phytohormones in the SAM, likely through regulation of nitrogen response and cellular redox status. Our data suggest that targeted phytohormone crosstalk engineering can be used to achieve increased reproductive growth and pathogen resistance.

**SIGNIFICANCE STATEMENT:** Plants constantly integrate environmental stimuli with developmental programs to optimize their growth and fitness. Excessive activation of the plant immune system often leads to decreased plant growth, a process known as the growth-defense tradeoff. Here, we adapted phytohormone levels in Arabidopsis reproductive tissues of autoimmunity mutants to change phytohormonal crosstalk and diminish the growth tradeoff, resulting in increased broad resistance to pathogens and decreased growth suppression. Similar approaches to phytohormone crosstalk engineering could be used in different contexts to achieve outcomes of higher plant stress resilience and yield.

## INTRODUCTION

Plants experience a variety of selective pressures depending on prevailing environmental conditions, resource availability, and physiological confines. Phytohormones and their associated networks integrate external and internal signals for refined plant cell decision-making during plant life, in a process known as phytohormonal crosstalk (1). The complexity of crosstalk amongst the different phytohormonal networks provides the signaling plasticity necessary to adapt to these constraints (2).

To ensure resilience, plants rely on tradeoffs as coordinated tactics to mitigate harm caused by sub-optimal conditions and overcome resource limitations. Of the several known plant tradeoffs, the most prominent is the growth-defense tradeoff, in which there is a cost to growth associated with the activation of defense responses to pathogens and pests (3, 4). This evolutionarily conserved plant survival strategy is complex and requires tight regulation of cellular signaling systems. While the canonical model of the growth-defense tradeoff assumes these two processes are mutually antagonistic, this idea has been challenged (5), instead favoring a model in which growth and defense are in continual dialogue to ensure plasticity in a fluctuating environment. In agreement with this, activation of plant immunity alters more than growth, leading to changes in plant developmental programs, underscoring the interconnection between development and immunity (6).

The relationship between plant growth and development and plant immunity can be attributed to phytohormone crosstalk. When a microbe or pest is present, the plant immune system is strategically activated in waves. Actively defending plants respond with upregulation of inducible defenses, such as production of reactive oxygen species (ROS), rapid apoplast alkalinization and calcium efflux, and expression of defense markers, to halt pathogen attack. Depending on the type of attacking pathogen or pest, accumulation of the defense phytohormones salicylic acid (SA), jasmonic acid (JA), and ethylene play a crucial role in activating appropriate immune responses. Notably, plants with over-accumulation or over-signaling of SA or JA display growth suppression similar to that observed during the growth-defense tradeoff due to activation of immunity (7–10), with the effects of shoot growth suppression being most studied. Shoot vegetative and reproductive growth are agronomically important traits determined at the shoot apical meristem (SAM). SAM activity is regulated by phytohormone signaling networks, namely, cytokinin (CK) and auxin (11). While SA has been shown to influence root apical meristem (RAM) regulation in a concentration-dependent manner, where low concentrations of SA applied to seedlings promote adventitious root growth and high concentrations halt RAM development (12), a clear association between defense hormones and SAM development has not been identified.

Amongst the known crosstalk interactions between phytohormones, the relationship between CK and SA may be of particular interest in relation to the growth-defense tradeoff evolutionary survival strategy. Although CK is known for its role in the regulation of cell division at the meristem level and overall shoot growth, it has also been shown to play an important role in the activation of defense, through cooperative interaction with the defense phytohormone SA (13–17). In contrast, it has been proposed that high SA content and/or signaling have inhibitory effects on the CK pathway, resulting in reduced growth (13). This suggests that these phytohormones may be at the crux of the signaling strategies plants use to maintain a balance between adequate defense activation and growth and development.

Phytohormones and their associated transcriptional hubs have been proposed as potential targets for plant bioengineering (6, 18, 19). Engineering of plant phytohormone crosstalk has been previously suggested as a way to achieve desired phenotypic outcomes (20), but the pleiotropic nature of phytohormone action and the complexity of crosstalk networks can restrict such efforts. Here, we show engineering of phytohormone crosstalk for the re-establishment of phytohormone homeostasis in specific tissues, resulting in increased pathogen resistance and reduced suppression of reproductive growth, demonstrating that phytohormone crosstalk engineering for plant improvement is possible. We show that the manipulation of the CK pathway in scenarios of constitutive immune activation leads to perturbation of the growth-defense tradeoff, conferring mutant plants with high broad-spectrum pathogen resistance and reproductive yield. In addition, the unique phytohormone balance of our genetic combinations suggests that plant defense phytohormones may function in the SAM and that deviations in nitrogen signaling and redox status likely regulate the growth-defense tradeoff. By furthering investigations of crosstalk phytohormone engineering, future efforts in synthetic biology can be used to develop advanced crops with increased pathogen resistance, superior plant yield, and abiotic stress resistance.

## RESULTS

### Defense-related phytohormones salicylic acid and jasmonic acid inhibit cytokinin signaling

Plants with increased levels of SA or JA content or signaling often display reduced plant growth (7–10). Previous work demonstrated that Arabidopsis mutants with decreased SA content showed increased sensitivity to CK, suggesting that SA has an inhibitory effect on CK signaling (13). Given the importance of CK to plant growth, we hypothesized that a negative interaction of the defense phytohormones SA or JA on CK signaling could be partially responsible for the growth tradeoffs usually associated with immune activation.

To understand the interaction between defense phytohormones and CK on plant growth, we analyzed publicly available Arabidopsis gene expression data for changes in the expression of cytokinin-regulated genes after SA and JA treatments. Three hours after application of 10 µM SA, most of the genes investigated were markedly downregulated (Figure 1A), suggesting that exogenous application of SA to Arabidopsis plants has an inhibitory effect on the expression of genes in the CK signaling and metabolic pathways. A similar, but less marked response, was observed in plants three hours after treatment with 10 µM JA (Figure 1B). Given the pronounced effect of SA in repressing CK-regulated gene expression, we further addressed the role of SA-dependent autoimmunity in the inhibition of CK signaling. The autoimmune mutants *constitutive expressor of PR genes 1* (*cpr1*), *cpr5*, and *suppressor of npr1-1 constitutive 1* (*snc1*) have elevated levels of SA and constitutive defense activation (10, 21). *cpr1*, *cpr5*, and *snc1* were examined for their levels of CK-regulated transcripts of *ARABIDOPSIS RESPONSE REGULATOR 7* (*ARR7*), *CYTOKININ OXIDASE/DEHYDROGENASE 4* (*CKX4*), *CYTOKININ RESPONSE FACTOR 6* (*CRF6*), and *EXPANSIN 1* (*EXP1*) by qRT-PCR. Results in Figure 1C show that, for the most part, these mutants displayed a general inhibition of CK-regulated gene expression. To further evaluate CK-regulated gene responsiveness to SA, we performed time course phytohormone induction assays on Arabidopsis seedlings treated with 100 µM SA or a 0.1% DMSO vehicle control, over the course of 24 hr. Both *ARR7* and *CRF6* were downregulated in the presence of SA at each timepoint, beginning 15 minutes after SA treatment (Figure 1D). Similarly, *EXP1* was downregulated by SA treatment at most of the timepoints. These data recapitulate results obtained from publicly available data (Figure 1A) and demonstrate that the suppression of CK-regulated gene expression by SA occurs rapidly upon phytohormone perception.

**Figure 1:**
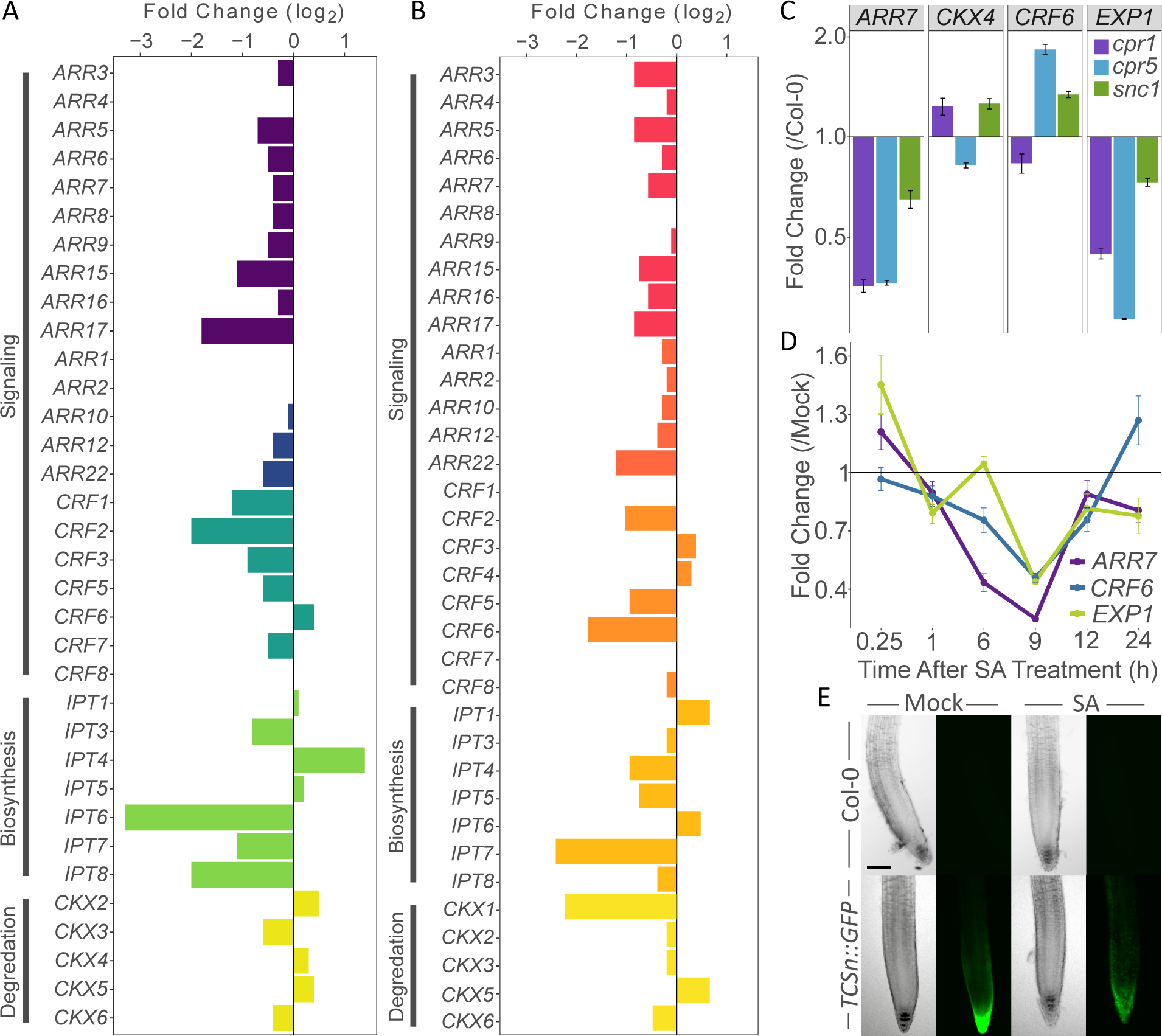
Defense phytohormones have an inhibitory effect on cytokinin signaling. Publicly available gene expression data of CK-regulated genes after (**A**) SA and (**B**) JA treatment. Six-day-old wildtype seedlings grown on plates were treated with 10 µM SA and tissue was collected 3 h after treatment. Expression values are log_2_-transformed and relative to the mock treatment. Data were acquired and analyzed using the e-northern tool of the Bio-Analytic Resource for Arabidopsis Functional Genomics [http://bar.utoronto.ca; (83)]. (**C**) CK-regulated gene expression of untreated mutants with increased SA biosynthesis/signaling *constitutive expressor of PR genes 1* (*cpr1*), *constitutive expressor of PR genes 5* (*cpr5*), and *suppressor of npr1-1 constitutive 1* (*snc1*). RNA was extracted from untreated 3-week-old leaf tissue. *ARABIDOPSIS RESPONSE REGULATOR 7* (*ARR7*), *CYTOKININ OXIDASE 4* (*CKX4*), *CYTOKININ RESPONSE FACTOR 6* (*CRF6*), and *EXPANSIN 1* (*EXP1)* transcript levels were determined by qRT-PCR relative to wildtype plants. Error bars represent s.e.m. from three technical replicates and correspond to upper and lower limits of 95% confidence intervals. For better visualization, the axis is log_2_-transformed. Data from one representative independent biological replicate are shown. At least three independent biological replicates of the experiment were conducted with similar results. (**D**) CK-regulated gene expression after 100 µM SA treatment over time. Wildtype seedlings were grown on 1X MS plates for 14 days. Seedlings were transferred to liquid MS and allowed to acclimate for 1 h on a shaker at 75 rpm under 120-150 µE light. SA, or solvent DMSO at 0.1%, was added to liquid culture. RNA was extracted from tissue collected at 0.25, 1, 6, 9, 12, and 24 h after phytohormone/mock treatment. Levels of *ARR7*, *CRF6*, and *EXP1* transcripts were determined by qRT-PCR relative to DMSO treatment. Error bars represent s.e.m. from three biological replicates and correspond to upper and lower limits of 95% confidence intervals. Data shown are three independent biological replicates pooled. (**E**) Fluorescence microscopy of 10 to 14-day-old Col-0 and transgenic *TCSn::GFP* roots treated with either mock or 50 µM SA. Representative images shown. Three independent biological replicates of the experiment were conducted with similar results. Brightfield (left); fluorescent imaging (right); n ≥ 20; scale bar = 100 µm.

We then investigated whether this rapid suppression of CK signaling by SA treatment could be maintained over time, or whether it was relieved shortly after stimulus perception. To address this question we used *TCSn::GFP* transgenic plants, harboring a synthetic transcriptional reporter for CK signaling (22), grown on Murashige and Skoog media (MS) plates supplemented with 50 µM SA or a mock control for 14 days. Mock-treated *TCSn::GFP* plants showed intense accumulation of GFP localized at the root tip (Figure 1E), as expected due to localized CK signaling (22, 23). When compared to the mock-treated plants, roots treated with SA displayed reduced GFP intensity, showing that SA can quickly repress CK-regulated gene transcription and that this suppression is sustained over extended periods of time, as long as the SA stimulus is present. Taken together, these experiments show that in addition to a positive crosstalk between CK and SA during plant immunity, a negative crosstalk of SA on CK responses also exists.

### Phytohormone crosstalk engineering

With the above-described phytohormone interactions in mind, we hypothesized that the disruption of the negative crosstalk of SA on CK signaling during active states of immunity could influence plant growth, relieving the growth-defense tradeoff. Given that the molecular mechanisms of this negative crosstalk are currently unknown, we addressed the relief of this negative crosstalk by engineering changes in phytohormone quantity, rather than signaling. Cytokinin oxidases/dehydrogenases (CKX) mediate the cleavage of the isoprenoid side chains of active CKs and their nucleosides, rendering the resulting CK species biologically inactive (24). In Arabidopsis, CKXs are encoded by a gene family of seven members, and loss-of-function mutations in *CKX* genes result in increased CK content, due to decreased degradation of CKs (24–26). We, therefore, hypothesized that reducing CKX activity in backgrounds with high levels of SA could render plants with increased cytokinin content, interfering with the negative crosstalk of SA on CK signaling, and relieving the suppression of growth due to immunity activation.

As fitness is arguably the most important agronomic trait, we aimed to manipulate the effects of the tradeoff on fruit and seed production. In order to engineer the crosstalk focusing on reproductive growth, we first addressed the tissue specificity and developmental regulation of *CKX* gene expression in Arabidopsis, by analysis of publicly available gene expression data. Analysis of the gene expression patterns of *CKX* genes revealed that *CKX3* and *CKX5* are predominantly expressed in reproductive tissues (Supplemental Figure 1), corroborating previous genetic work (25). *suppressor of npr1-1 constitutive 1* (*snc1-1*) is an autoimmune mutant, and this allele harbors a gain-of-function mutation in a gene encoding a nucleotide-binding leucine-rich repeat (NLR) protein, leading to constitutive defense activation, high SA content, dwarfed morphology, and reduced fruit and seed production (10). To engineer the crosstalk with the goal of diminishing the reproductive tradeoff, we crossed *snc1* and *ckx3,5* plants and obtained a triple mutant, which we named *s35*.

### *s35* displays a novel reproductive growth phenotype

To determine the effect of changing the phytohormone crosstalk between CK and SA on plant growth, we characterized the *s35* triple mutant, as well as its parentals and wildtype Col-0 plants, with regard to several aspects of plant growth. When grown in short days, which promotes vegetative growth, *s35* plants were slightly, although not significantly, larger than *snc1* plants, but still exhibited a smaller rosette than wildtype (Figure 2A and 2B). To evaluate root growth, plants of all genotypes were grown on vertical MS plates. The *snc1* mutant had shorter roots, and this phenotype was maintained in *s35* plants (Supplemental Figure 2A and 2B), however RAM cell count was not significantly changed in all genotypes (Supplemental Figure 2D). In addition, the roots of *ckx3,5* plants contained additional layers of root cap, and the same phenotype was observed in *s35* plants (Supplemental Figure 2C). Similarly, an assay for CK-induced chlorophyll retention of leaves during dark-induced senescence showed that retained levels of chlorophyll in *s35* plants were not higher than in *snc1* plants (Supplemental Figure 3A), and a primary root growth assay showed that root growth of all genotypes was similarly affected by CK (Supplemental Figure 3B). Taken together, these results show that the combination of the *ckx3,5* mutations with *snc1* in the *s35* plants does not lead to large phenotypic changes in vegetative tissues.

**Figure 2:**
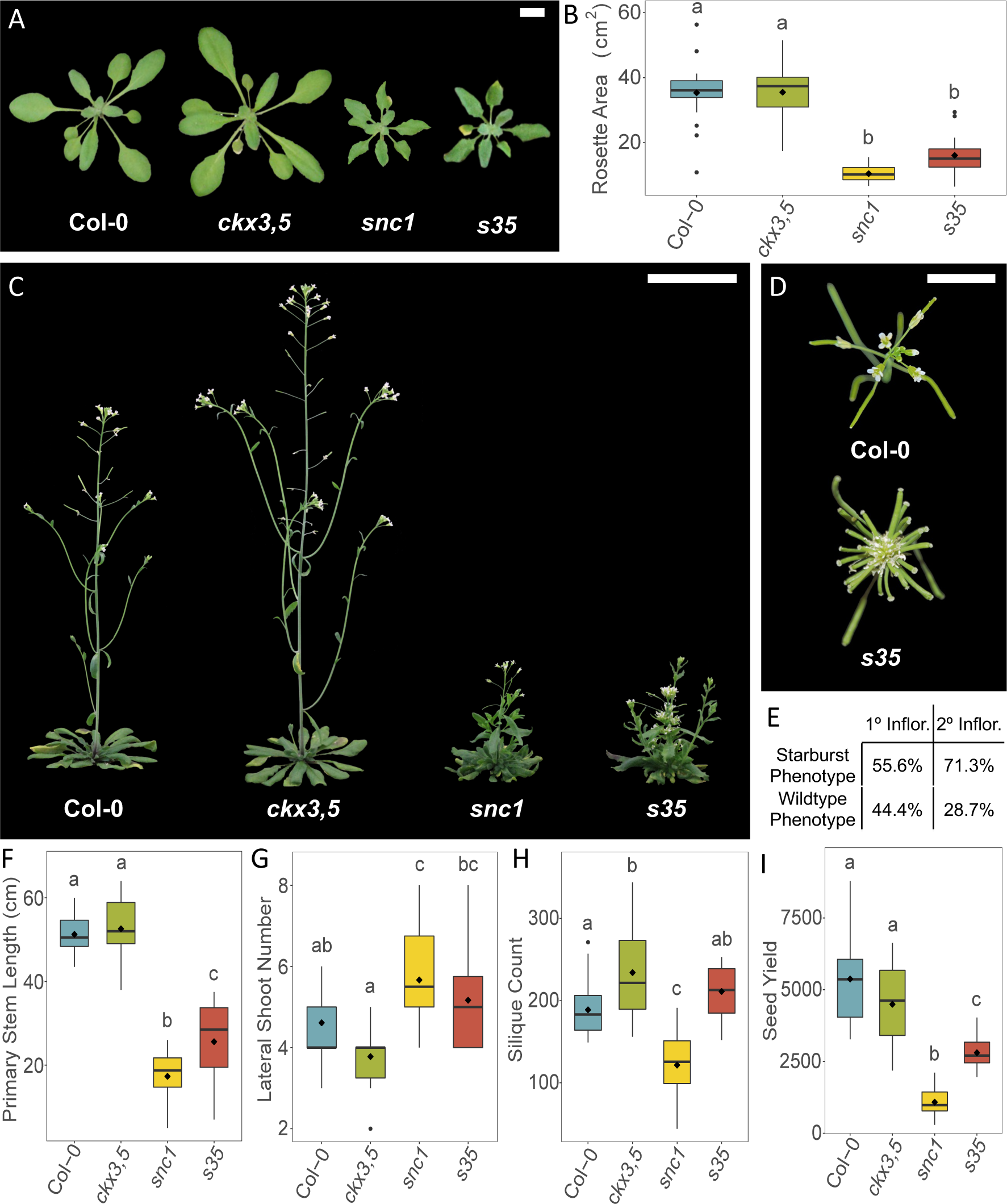
The *s35* triple mutant has a novel reproductive growth phenotype. (**A**) Representative vegetative growth phenotypes of each genotype 30 days after germination. Scale bar = 1 cm. (**B**) Rosette area of all genotypes 30 days after germination. (**C**) Representative flowering phenotypes of 50-day-old plants. Scale bar = 5 cm. (**D**) Representative inflorescences between wildtype Col-0 and *s35* at 8 weeks old. *s35* has a novel “starburst” inflorescence phenotype. Scale bar = 1 cm. (**E**) Penetrance of novel starburst phenotype of *s35* primary and secondary inflorescences. n = 18. (**F**) Stem length of the primary stem, (**G**) number of lateral shoots, and (**H**) average silique count of whole plants 10 weeks after germination. (**I**) Whole plant seed yield collected 12 weeks after germination. For (**B**), (**F**), (**G**), (**H**), and (**I**), means labeled (♦), and letters indicate differences at a P < 0.05 significance level using a one-way ANOVA analysis with a Tukey post-hoc correction. n = 18.

Given our emphasis on tradeoffs of reproductive growth, *s35* plants were assessed for any growth penalties in reproductive tissues (Figure 2C to 2I). *s35* plants retained the reduced apical dominance phenotype characteristic of *snc1* plants (10), with reduced primary stem growth (Figure 2F) and more than double the number of secondary lateral shoots compared to wildtype (Figure 2G). At the terminal end of shoots, *s35* plants displayed a novel inflorescence growth pattern. The siliques were compressed as the internodes between successive organs were drastically reduced. This results in a distinctive inflorescence growth pattern at the shoot apex we coined the “starburst” phenotype (Figure 2D). The starburst phenotype has an altered penetrance in primary shoots versus secondary shoots, with about 55% of primary shoots and about 71% of secondary shoots displaying the starburst phenotype (Figure 2E). *ckx3,5* had a significantly increased number of siliques when compared to wildtype (Figure 2H), while *snc1* plants had a significant reduction in silique number, as previously described in these backgrounds (10, 25). In contrast, the silique count of *s35* was similar to that of Col-0, suggesting that an increase in CK content could rescue the reduced silique production observed in the *snc1* background. In addition to increased silique production, *s35* had over double (almost 260%) the seed yield compared to *snc1* (Figure 2I), although not the same seed yield as wildtype plants. Examination of siliques revealed a higher number of aborted seeds or unfertilized ovules in *s35* plants as compared to Col-0 (data not shown), which may contribute to the reduced seed production in this genotype (Figure 2I).

### Increased reproductive growth of *s35* results from re-established CK levels and altered SAM function

To determine if the starburst phenotype of the *s35* mutant was the result of alterations in the SAM, scanning electron microscopy (SEM) of SAMs was conducted (Figure 3A to 3E). *ckx3,5* displayed increased meristem diameter and number of floral organ primordia compared to Col-0, as previously described (Figure 3F) (25). The *snc1* mutant had a reduced meristem size compared to Col-0, implicating SA as a negative regulator of meristem size. Most notably, *s35* meristems often had remarkably altered meristematic growth: some meristems displayed the wildtype phenotype (Figure 3D), while others had severely compacted floral organ primordia of many stages (27), swollen sepals, and deviation from the typical SAM phyllotactic pattern (Figure 3E). As the plant matures, this irregular meristematic patterning develops into the distinctive starburst inflorescence phenotype of *s35* plants (Figure 2D).

**Figure 3:**
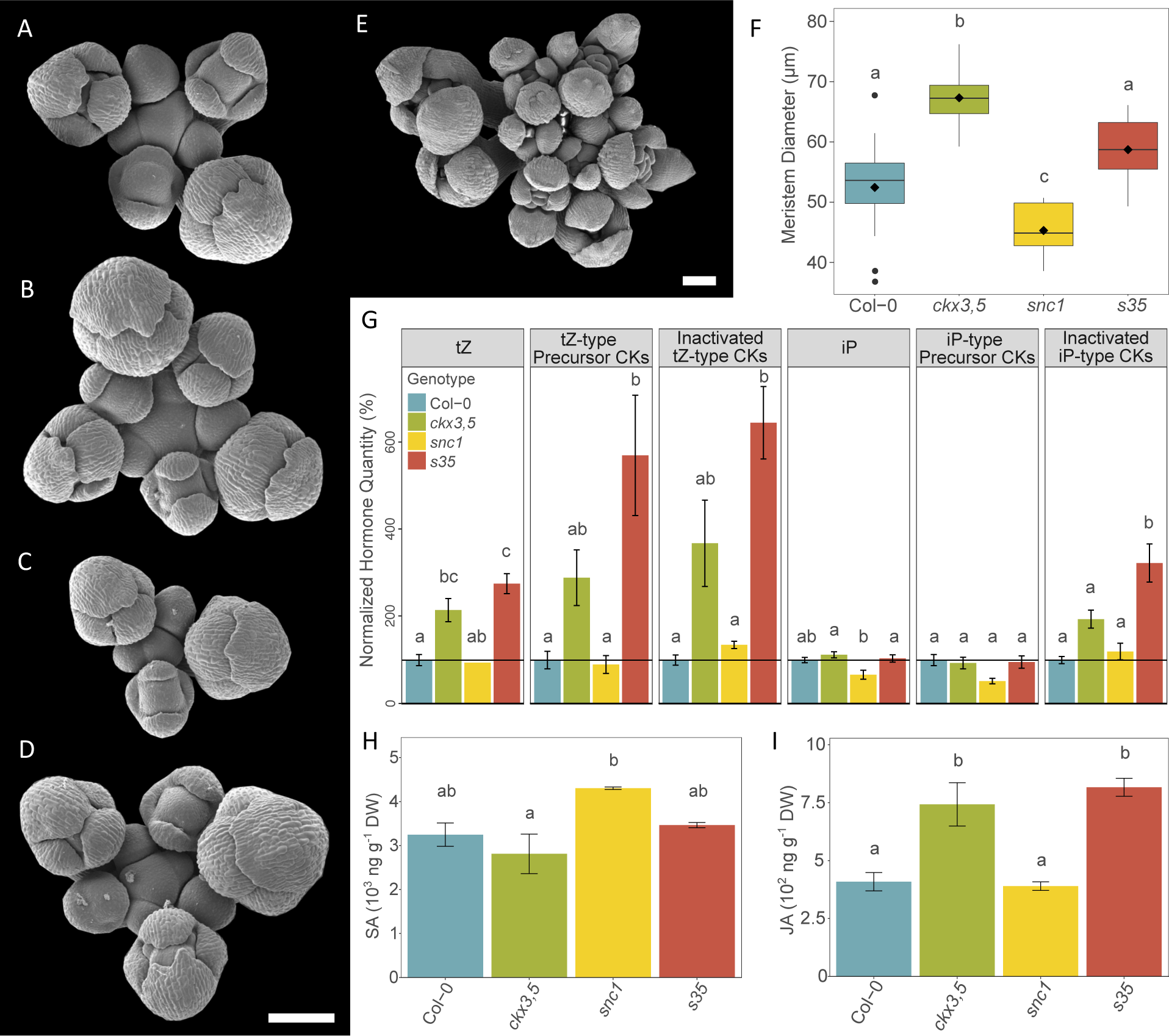
The *s35* triple mutant has altered inflorescence development due to changes in phytohormone balance. Representative electron scanning micrographs of hand-dissected primary shoot apical meristems of (**A**) Col-0, (**B**) *ckx3,5*, (**C**) *snc1*, and (**D**), (**E**) *s35*. Scale bars = 100 µm. (**F**) Diameter of hand-dissected primary shoot apical meristems. Means labeled (♦), and letters indicate differences at a P < 0.05 significance level using a one-way ANOVA analysis with a Tukey post-hoc correction. n ≥ 12. Basal phytohormone quantification of different CK species normalized to Col-0: (**G**) tZ, tZ-type CK precursor, sum of tZR and tZRPs, inactivated tZ-type CK, sum of tZ7G, tZZ9G, tZOG, tZROG, and tZRPsOG, iP, iP-type CK precursor, sum of iPR and iPRPs, and inactivated iP-type CK, sum of iP7G and iP9G. See Supplemental Table 1 for absolute values. (**H**) SA and (**I**) JA quantification of untreated inflorescence tissue. For (**G**), values represent percentage of wildtype CK species amount ± the percentage s.e.m. of three technical replicates. For (**H**) and (**I**), values represent the mean ± the s.e.m. of three technical replicates. For (**G**), (**H**), and (**I**), each replicate contained ≥ 30 inflorescences. Letters indicate differences at a P < 0.05 significance level using a one-way ANOVA analysis with a Tukey post-hoc correction.

To demonstrate that the phenotype of *s35* plants was associated with increased CK content, CK species were quantified in inflorescence tissue by mass spectrometry (see Supplemental Table 1 for absolute quantities of all species measured). The main bioactive forms of CK are considered to be *trans*-Zeatin (tZ) and N^6^-(Δ^2^-isopentenyl)adenine (iP), which play differing roles (28). tZ-type precursor CKs (most often tZ-riboside) are thought to travel from the root to the shoot in xylem tissue, during nutrient sensing (29, 30), while iP-type CKs move from shoot to root in the phloem. Quantification of CK species in the *ckx3,5* reproductive tissues showed that *ckx3,5* plants had higher levels of tZ and tZ-type CKs, both precursor and inactive species, compared to wildtype (Figure 3G). Similarly, *s35* also displayed increased levels of tZ-type CKs. iP-type CKs were at relatively low levels compared to tZ-types in all genotypes and particularly in *snc1* plants, although *s35* had significantly higher levels of inactivated iP-type species compared to all genotypes, suggesting possible modifications to CK metabolism or translocation in the triple mutant.

To determine whether an increase in CK was sufficient to increase reproductive growth in immunity-activated backgrounds and produce the starburst phenotype, *snc1* plants were sprayed with either a 1 µM or 10 µM solution of the synthetic CK 6-benzylaminopurine (BA), or a mock control, two times a week for three weeks after bolting. BA-treated *snc1* inflorescences showed a higher number of siliques and shortened internodes that phenocopied the starburst phenotype in a dose-dependent manner (Supplemental Figure 4). Further, to address whether *ckx3,5* mutations could also rescue suppression of growth in other autoimmunity mutants, we crossed *ckx3,5* to *cpr1*, an autoimmunity mutant that also displays suppression of vegetative and reproductive growth. *cpr1 ckx3,5* triple mutant plants (named *c35*) had inflorescences that resembled those of *s35*, with starburst phenotypes (Supplemental Figure 5). Thus, this altered reproductive phenotype is likely due to the combinatorial effects of the *ckx3,5* mutations in the context of autoimmunity and effective in other autoimmune backgrounds.

The results above led us to ask the question of whether the levels of defense phytohormones, like SA and JA, were changed in the *s35* mutant. Free SA and JA were quantified in inflorescence tissue dissected to young, unopened flower buds (stage 15) (27). *snc1* inflorescences had the highest accumulation of SA (Figure 3H), an outcome expected given its autoimmune background. If SA acts to repress SAM activity, either directly or indirectly, this disparity in SA levels could underlie the dwarfed versus enlarged SAM phenotypes observed in *snc1* and *ckx3,5*, respectively (Figure 3B and 3C). Moreover, Col-0 and *s35* had intermediate SA content, similar to their intermediate SAM phenotypes (Figure 3A and 3D). JA levels were higher in the *ckx3,5* and *s35* inflorescences, suggesting that CK promotes JA biosynthesis in reproductive tissues. The novel combination of autoimmunity, resulting in increased SA content, and CK overabundance in these tissues, likely contributes to the *s35* starburst phenotype.

### *s35* displays higher resistance to biotrophic and necrotrophic pathogens due to changes in phytohormone crosstalk

*snc1* plants have increased resistance to biotrophic pathogens, such as *Pseudomonas syringae* pv. *tomato* DC3000 (*Pst*) and *Hyaloperonospora arabidopsidis* Noco2 (*Hpa*) (10). We wanted to determine whether the partial rescue of *snc1* reproductive growth by the *ckx3,5* mutations altered the *snc1* defense phenotype and caused a shift in the growth-defense tradeoff. After inoculation with *Hpa*, *s35* retained the resistance observed in *snc1* plants, allowing for a low number of sporangiophore production and reduced hyphal growth than wildtype plants (Figure 4A). A similar response of decreased susceptibility in *s35* plants was observed after inoculation with *Pst* (Figure 4B). A characteristic of the *snc1* mutant is the constitutive activation of defense genes, such as *PATHOGENESIS-RELATED 1* (*PR1*) (10). Like *snc1*, *s35* plants showed constitutive activation of *PR1* (Supplemental Figure 6). *ckx3,5* plants also displayed constitutive *PR1* expression, reflecting the positive crosstalk of increased CK on SA (13). When challenged with *Pst*, *snc1* and *s35* showed even higher levels of *PR1* expression when compared to mock treatment, with *s35* plants showing the highest induction (Figure 4C). *snc1* and *s35* had elevated levels of free SA in leaves, reflecting their autoimmune nature (Figure 4D).

**Figure 4:**
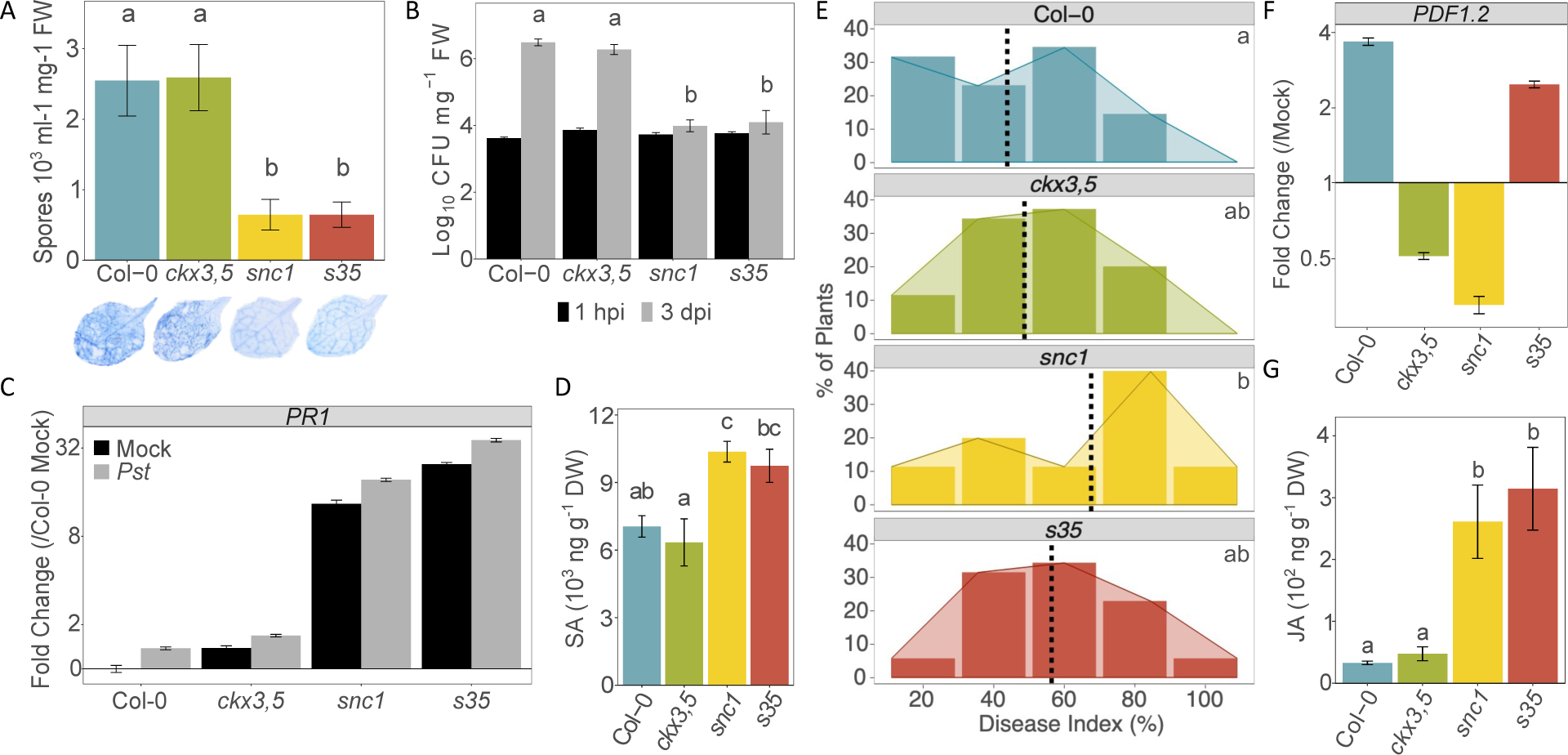
The *s35* triple mutant displays resistance against biotrophic pathogens and reduced susceptibility to a necrotrophic pathogen. (**A**) *Hyaloperonospora arabidopsidis* Noco2 (*Hpa*) spore counts of each genotype at 3 dpi. Data shown are three independent biological replicates pooled. Below, representative leaves after trypan blue straining of *Hpa* hyphal/oospore growth. (**B**) Growth of *Pseudomonas syringae* pv. *tomato* DC3000 (*Pst*) at day 0 (T0) and at 3 dpi (T3). Data from one representative independent biological replicate are shown. At least three independent biological replicates of the experiment were conducted with similar results. (**C**) *PATHOGENESIS-RELATED 1* (*PR1*) expression 1 h after *Pst* inoculation (OD = 0.05). Levels of the indicated transcripts were determined by qRT-PCR relative to mock-treated wildtype plants and normalized to *UBIQUITIN 10* (*UBQ10*). Error bars represent s.e.m. from three biological replicates and correspond to upper and lower limits of 95% confidence intervals. For better visualization, the axis is log_2_-transformed. Data pooled from three independent biological replicates shown. (**D**) SA quantification of untreated 6-week-old leaf tissue. Values represent the mean ± the s.e.m. of three technical replicates. (**E**) Histograms of qualitative disease index of 6-week-old plants after *Botrytis cinerea* spray inoculation (0.5×10^4^ spores/mL). Photographs of individual inoculated plants were taken at 96 hpi. Plants were categorized based on disease severity, where disease index represents percentage of rosette covered in necrotic lesions. Four independent biological replicates were conducted with similar results. Data from all four independent biological replicates were pooled. Dotted lines indicate average disease index per genotype. Letters indicate differences of disease index averages at a P < 0.05 significance level using a two-way ANOVA analysis with biological replicate as a blocking factor. (**F**) JA quantification of untreated 6-week-old leaf tissue. Values represent the mean ± the s.e.m. of three technical replicates. (**G**) *PLANT DEFENSIN 1.2* (*PDF1.2*) gene expression 6 h after *B. cinerea* spray inoculation (0.5×10^4^ spores/mL). Levels of the indicated transcripts were determined by qRT-PCR normalized to *UBIQUITIN 10* (*UBQ10*) and relative to mock-treated samples. Error bars represent s.e.m. from three technical replicates and correspond to upper and lower limits of 95% confidence intervals. At least three independent biological replicates of the experiment were conducted with similar results. Data from one representative independent biological replicate are shown. For better visualization, the axis is log_2_-transformed. For (**A**), (**B**), (**D**), and (**F**), letters indicate differences at a P < 0.05 significance level using a one-way ANOVA analysis with a Tukey post-hoc correction.

Increased resistance to biotrophic pathogens is usually accompanied by increased susceptibility to necrotrophs, due to the antagonistic interaction between the SA and JA pathways (31). The increased resistance to biotrophs observed in the *s35* triple mutant prompted us to determine its susceptibility to necrotrophs. Wildtype, *ckx3,5*, *snc1*, and *s35* plants grown under short-day conditions were inoculated with the necrotrophic fungus *Botrytis cinerea* by spray inoculation and susceptibility determined by percentage of rosette area with necrotic lesions in each genotype (Figure 4E). Wildtype plants displayed an average of 46% of the rosette area with necrotic lesions, with very few plants showing over 80% of necrotic rosette area. *snc1*, on the other hand, had a higher percentage of plants with over 80% of necrotic rosette area, and a shift of 18% in susceptibility over wildtype plants. To our surprise, *s35* plants, despite being resistant to biotrophic pathogens (Figures 4A and 4B), were less susceptible than *snc1* plants, with a shift of only 8% in susceptibility over wildtype plants, comparable to *ckx3,5* plants (Figure 4E).

Quantification of JA levels in all genotypes showed an enhanced accumulation of free JA in both *snc1* and *s35* (Figure 4F), indicating that the constitutive activation of an NLR protein by the *snc1-1* gain-of-function mutation increases resistance to pathogens by activating both the SA and JA pathways. *PLANT DEFENSIN 1.2* (*PDF1.2*) is a widely used marker gene for defense against necrotrophic pathogen attack and JA signaling. As expected, after *B. cinerea* inoculation *PDF1.2* was induced in wildtype plants. This induction was not seen in *ckx3,5* and *snc1*, likely due to their increased SA signaling leading to suppression of the JA pathway, however, *PDF1.2* expression levels were higher after pathogen treatment in *s35* (Figure 4G). The increased JA content but reduced *PDF1.2* expression of *snc1* and *s35* plants indicates that the suppression of JA responses by SA happens at the signaling level, and not at the level of phytohormone content. Moreover, this suppression on the JA pathway is reduced in *s35* plants, resulting in increased resistance to both biotrophs and necrotrophs in this genetic combination.

### Altered ROS compartmentalization in plant tissues is associated with increased reproductive yield and resistance to pathogens

To gain insight into the biological processes occurring within the reproductive tissue to yield the novel *s35* phenotype, we performed RNA sequencing analyses of inflorescences across genotypes dissected to stage 13+ (27, 32). The Col-0 transcriptome was used as a control to identify differentially expressed genes (DEG) in *ck3,5*, *snc1,* and *s35* plants. At a threshold of Benjamini-Hochberg’s adjusted P value < 0.05, our analysis revealed 648 DEG (172 up and 476 down) in *ckx3,5*, 5767 DEG (3145 up and 2622 down) in *snc1*, and 4892 DEG (2575 up and 2317 down) in *s35* (Figure 5A). The high number of DEG in the *snc1* and *s35* backgrounds in comparison to the *ckx3,5* double mutant underscores the strong influence of the *snc1* gain-of-function mutation on the global inflorescence transcriptome, due to the activation of several immune-related processes. DEG, at a log_2_ fold change (FC) cutoff of 1, were then analyzed between the three genotypes in comparison to wildtype plants (Figure 5B, 5C, and 5D), and gene ontology (GO) analysis was used to characterize the DEG from the gene comparison lists into biological process enrichment categories via the PANTHER 17.0 database (33). The GO terms relevant to the *ckx3,5* mutant mainly showed alterations to reproductive processes, most related to pollen tube growth. The GO terms “regulation of developmental growth” (Fold Enrichment (FE) = 14.34; False Discovery Rate (FDR) = 1.43E-05) and “regulation of reproductive process” (FE = 6.64; FDR = 2.93E-03) were shared between *ckx3,5* and *s35* (data not shown), potentially pointing to the increased SAM activity in these mutants. As expected, *snc1* upregulated DEG were largely categorized into defense-related GO terms (Figure 5E), many of which were shared with *s35* (Figure 5F). Interestingly, the GO term with the highest fold enrichment unique to the *snc1* background was “negative regulation of cytokinin-activated signaling pathway” (FE = 52.8; FDR = 0.0121), providing further evidence of the inhibitory effect of SA on CK signaling (as seen in Figure 1).

**Figure 5:**
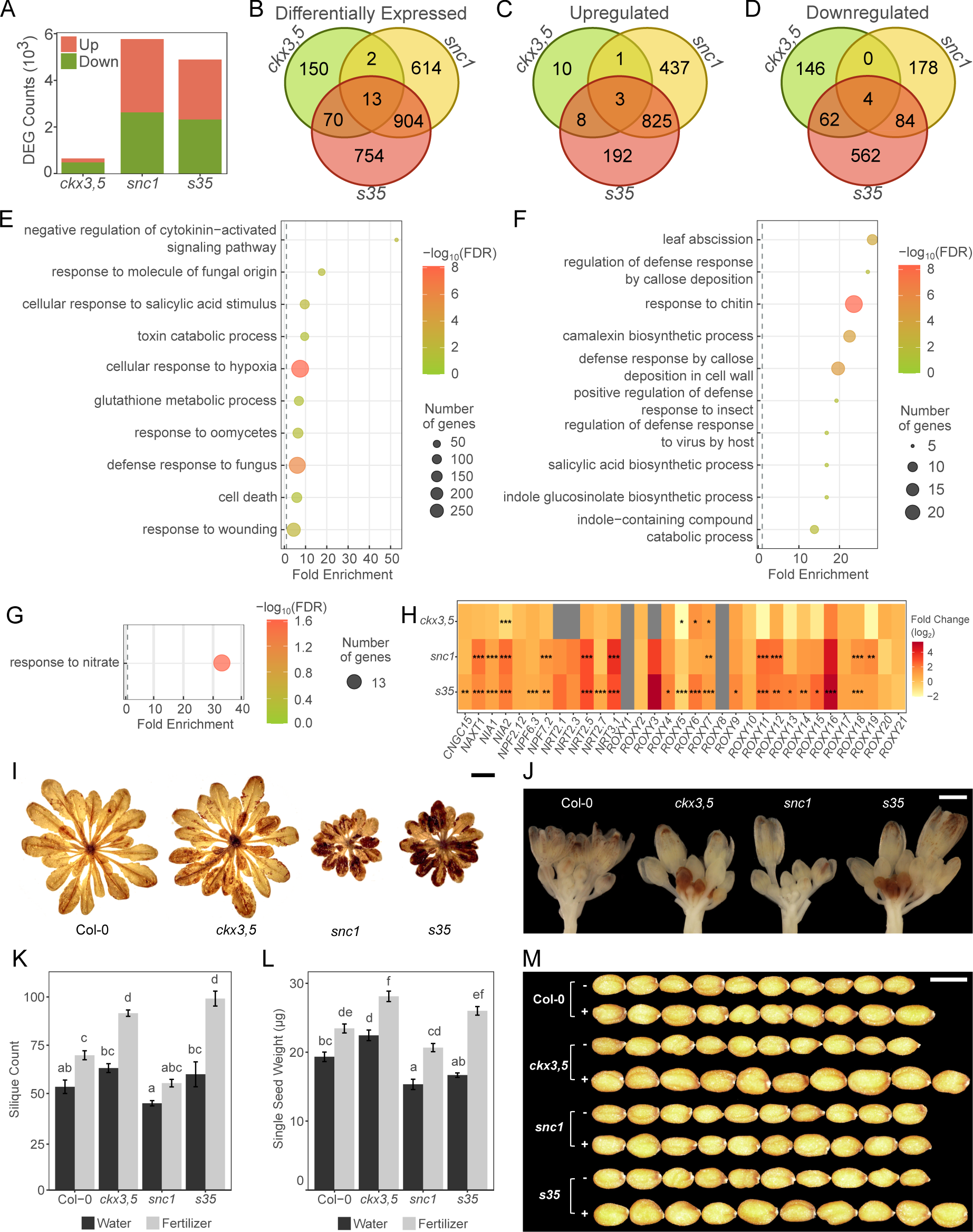
Transcriptome analysis of inflorescences reveals that *s35* inflorescences have a unique *ROXY* gene family expression pattern and could mediate reproductive growth in response to fertilization. (**A**) Number of DEG (up- and downregulated) of each genotype compared to the wildtype transcriptome of untreated inflorescence tissue. For the three biological replicates per genotype, each sample was collected developmentally, when the primary shoot had at least one fully expanded flower present (stage 13+). DEG lists were compared amongst the mutant genotypes using Venn diagrams: (**B**) all differentially expressed genes, (**C**) upregulated genes, and (**D**) downregulated genes. An arbitrary log_2_ fold change cutoff of 1 was used to reduce the DEG lists before comparison and in analyses hereafter. (**E**) Gene Ontology (GO) term enrichment of uniquely upregulated DEG in the *snc1* background [437 genes from (**C**)]. (**F**) GO term enrichment upregulated DEG shared between *snc1* and *s35* [825 genes from (**C**)]. (**G**) GO term enrichment of uniquely upregulated DEG in *s35* [192 genes from (**C**)]. Terms with a significant fold enrichment of FE ≥ 12 shown. (**H**) Gene expression from the transcriptomic analysis of nitrogen-related genes and the CC-type glutaredoxin (*ROXY*) gene family in untreated inflorescences. The gene list was assembled from relevant literature. Statistically significant differences from wildtype inflorescences (FDR) are represented by asterisks (* = P value < 0.05, ** = P value < 0.01, *** = P value < 0.001). (**I**) Representative 6-week-old rosettes and (**J**) inflorescence meristems with hydrogen peroxide (H_2_O_2_) stained with 3, 3-diaminobenzidine (DAB). Col-0, *ckx3,5*, *snc1*, and *s35* were watered with or without 1X Miracle-Gro Water Soluble All Purpose Plant Food. At least two independent biological replicates of the experiment were conducted with similar results. (**K**) Average silique count of primary stems 12 weeks after germination. (**L**) Estimated single seed weight. The single seed weight was estimated from the weight of 200 seeds per plant. (**M**) Ten representative seeds of all four genotypes from plants watered with either water (-) or 1X Miracle-Gro Water Soluble All Purpose Plant Food (24-8-16) (+). For (**E**), (**F**), and (**G**), gene lists from the respective comparisons were analyzed for pathway enrichment using PANTHER 17.0 database [http://go.pantherdb.org/; (33)]. Only GO terms with an FDR-corrected P value < 0.05 were considered significantly enriched. For (**K**) and (**L**), letters indicate differences at a P < 0.05 significance level using a two-way ANOVA analysis with a Tukey post-hoc correction. n = 18. For (**I**), scale bars = 1 cm. For (**J**), scale bar = 100 µm. For (**M**), scale bar = 500 µm.

Amongst the 192 uniquely upregulated DEG in the *s35* background, only a single significant GO term was returned, “response to nitrate” (FE = 32.01; FDR = 0.0226; Figure 5G). Within this category were several genes encoding CC-type glutaredoxins (*ROXY*s or *GRX*s). Glutaredoxins are thiol oxidoreductases, that bind to glutathione (GSH), are involved in post-translational modifications of other proteins, and also participate in the control of ROS through scavenging (34). CC-type glutaredoxins are unique to plants, with 21 gene family members in Arabidopsis, and mutations in these genes have revealed roles for them in development, nitrogen starvation, and stress responses (35). A total of 11 *ROXY* genes were significantly upregulated in *s35* (Figure 5H). ROS signaling plays an important role in the regulation of plant growth, including meristematic activity in the SAM (36). We therefore addressed whether ROS levels could be linked to altered reproductive phenotypes observed in *s35* plants. A histochemical assay for H_2_O_2_ staining showed a high basal accumulation of H_2_O_2_ in leaf tissue *snc1* and *s35* (Figure 5I), a known phenotype associated with immunity activation (37–40). In inflorescence tissue, however, *snc1* plants accumulated less H_2_O_2_ than wildtype plants, and *ckx3,5* and *s35* had markedly pronounced staining in young flower buds near meristematic tissue (Figure 5J). Thus, H_2_O_2_ is spatially sequestered in vegetative tissues in the *snc1* background, and the presence of this reactive species near the SAM is associated with the increased reproductive growth of *ckx3,5* and *s35*.

### Fertilization can rescue seed yield in *s35* plants

ROXYs also play a role in nutrient sensing, specifically nitrate signal transduction (41). CK levels have a positive correlation with nitrate (29, 30) as CK is considered a long-range signal for nitrate availability (42). This role of ROXYs, along with the increased expression of genes encoding several nitrate transporters (*NRT*s and *NPF*s), nitrate reductase (*NIA1/2*), and nitrate response genes in *s35* plants (Figure 5H) led us to evaluate whether nitrate availability could induce the starburst phenotype in our parental mutants. Plants of all genotypes were fertilized at each watering with fertilizer solution containing a 24-8-16 nitrogen-phosphorus-potassium (NPK) macronutrient ratio. Heavy fertilization did not phenocopy the starburst in either *ckx3,5* or *snc1* (data not shown), yet all genotypes had improved growth and increased silique number (Figure 5K), particularly *s35* plants. Most importantly, under fertilization, the seeds of *s35* plants were bigger and heavier (Figure 5L and 5M), even in comparison to wildtype plants. Thus, by increasing seed size, fertilization can partially compensate for the reduced seed yield of the *s35* genotype in comparison to wildtype plants.

## DISCUSSION

As they grow, plants integrate a variety of environmental cues into developmental programs, to optimize their fitness in different environments. Activation of plant immunity by chemical or biological agents often results in suppression of plant growth, with consequences to plant fitness. This phenomenon, known as the growth-defense tradeoff, has been widely studied at the molecular, physiological, and ecological levels. This suppression of growth decreases overall plant yield and prevents the use of mutations leading to higher levels of immunity and disease protection in plant breeding (43, 44).

Cytokinins, first discovered for their role in cell division, control several cellular and physiological processes (45), including plant reproductive yield, through the regulation of SAM activity. In recent years, a positive crosstalk between CK and the defense phytohormones SA and JA was identified to increase plant immunity, in Arabidopsis and other plant species (13, 14, 16). Here, we demonstrate a previously suggested negative crosstalk action of JA, and most significantly SA, on CK signaling (13), and show that this crosstalk exists in contexts of constitutive immunity. Given that SA has been determined to account for the majority of growth suppression in the *snc1* background (10, 46, 47), and that *snc1* inflorescences do not display high levels of JA (Figure 3I) or a strong signature of altered JA signaling (Supplemental Figure 8), we hypothesized that modifying the crosstalk between the CK and SA pathways would relieve the suppression on CK signaling and restore reproductive growth tradeoffs caused by SA-based defense activation.

Phytohormonal crosstalk exists when the combinational signal of different pathways results in an outcome different from the signal of each pathway by itself (1). Crosstalk can occur at the level of signaling, by physical interaction followed by post-translational modifications of proteins from different phytohormone pathways, or at the level of phytohormone quantity, by changes of gene expression or activity of enzymes involved in other phytohormone’s biosynthesis and catabolism. Instead of targeting phytohormone signaling, we opted to re-engineer the CK-SA crosstalk via changing phytohormone levels. By targeting mutations in genes encoding CKXs, the enzymes responsible for CK degradation, we were able to increase CK content in Arabidopsis plants, while constitutively activating immunity due to either a gain-of-function mutation in a positive regulator of immunity (*snc1-1*) or loss-of-function mutation in a negative regulator of immunity (*cpr1*). Targeting *CKX* genes mostly expressed in reproductive organs (*CKX3* and *CKX5*) allowed us to change the crosstalk in specific tissues and largely avoid unwanted pleiotropic effects in other parts of the plant.

We observed significant morphological changes in the resulting *s35* mutant, particularly in reproductive tissues, with the reproductive cost of immunity characteristic of the *snc1* phenotype partially rescued by the *ckx3,5* mutations. Although *s35* mostly retained dwarfed vegetative growth, the number of siliques of *s35* plants was increased in comparison to both *snc1* and Col-0 plants, although seed yield was reduced in relation to Col-0. An increase in seed pods that is not similarly reflected in seed number has been previously observed in *ckx* mutants and was attributed to heterostyly that occurred in certain environmental conditions, which diminished self-fertilization (25). While we did not look for evidence of heterostyly in *s35* or *ckx3,5* plants under our conditions, we did observe that, like *s35*, *ckx3,5* plants had increased silique number but not an increased number of seeds.

We observed that the increased number of siliques of *s35* plants was related to changes in inflorescence structure, phyllotaxis, and SAM size, resulting in a phenotype we coined “starburst”. Normative growth patterning of shoot meristematic tissue is under tight regulation by phytohormone crosstalk, including cytokinins (11, 48–52). While a role for SA has been identified in RAM regulation in a concentration-dependent manner (12), a role for SA in the SAM has not been shown. Notably, we provide evidence for a link between endogenous SA levels and SAM development in the *snc1* background, which has high accumulation of SA in inflorescence tissue and smaller meristems, suggesting that SA antagonizes SAM growth. Moreover, *s35* not only has increased meristematic activity like *ckx3,5* but also altered silique phyllotaxy, likely by modification of CK-regulated functions in the SAM (53). To our knowledge, this is the first time that SA, or its downstream signaling, has been implicated in SAM developmental regulation. Further experimentation is necessary to identify the mechanisms behind this.

SA and ROS production are common outcomes of immunity activation, and ROS are also involved in the regulation of SAM function (36). In the case of *ckx3,5* and *s35* plants, increased H_2_O_2_ staining is observed in the young flower buds adjacent to meristematic tissue, and this staining is absent in *snc1* plants, in which H_2_O_2_ accumulation is restricted to vegetative tissue. While ROS accumulation is commonly associated with suppression of growth and activation of cell death pathways, it is also a necessary step for redox changes of proteins that trigger cell proliferation in both animal and plant stem cells (54–58), and too much antioxidant activity can in fact impair meristem function (59). Furthermore, the SAM is a hypoxic environment (60), and hypoxia is known to induce hydrogen peroxide production in rapidly dividing plant cells. The accumulation of H_2_O_2_ in *ckx3,5* and *s35* meristems suggests a positive role for this ROS species in the proliferation of cell division in meristematic tissue, likely connected with the enhanced cytokinin accumulation of these genetic backgrounds.

ROS-dependent control of the inflorescence meristem has been demonstrated, by mechanisms that include redox regulation of transcription factors and changes in the redox state of the stem cell niche (61) involving ROXYs. In maize, the ROXYs MALE STERILE CONVERTED ANTHER 1 (ZmMSCA1), ZmGRX2 and ZmGRX5 act redundantly in SAM development, targeting the FASCIATED EAR 4 (ZmFEA4) TGA transcription factor for post-translational modification, and influencing the redox state of the stem cell niche (61). *grx* double and triple knockouts have defects in ear development with significantly fewer kernels than wildtype and decreased SAM size (61). In Arabidopsis, SAM size is also tightly regulated by nitrate availability in the soil and associated with systemic CK signaling (62). A direct link between ROXYs, nitrogen sensing, CK signaling, and meristematic activity has been proposed in roots, mediated by ROXY6 and/or ROXY9, referred to in this work as CEP DOWNSTREAM 1 (CEPD1) and CEPD2, which act downstream of perception of the C-TERMINALLY ENCODED PEPTIDE (CEP) peptide hormone. CEPDs coordinate crosstalk between CEP and CK for ideal regulation of root growth in response to nitrogen acquisition (63). While we don’t demonstrate a role for ROXYs in the regulation of *s35* reproductive phenotypes, their altered gene expression, coupled with the effect of fertilization (including nitrogen) in the seed output, plus the spatial restriction of ROS observed in this genotype, highly suggest a function for these proteins in these processes.

Another result of our crosstalk modification was the increased resistance of *s35* plants to both biotrophic and necrotrophic pathogens. The *snc1-1* gain-of-function mutant used in this study is highly resistant against biotrophic pathogens, like *Pst* and *Hpa* (10), and these results were confirmed in our work. Resistance to biotrophic pathogens is usually accompanied by SA accumulation, which can antagonize defense responses to necrotrophs, mediated by JA (31, 64, 65), hindering broad-spectrum resistance. Yet, *s35* is less susceptible to the necrotrophic fungus *B. cinerea* when compared to *snc1*, and thus represents a mutant combination that provides higher resistance to both biotrophs and necrotrophs. Recently, a function for CKs in immunity against necrotrophic pathogens was observed in tomato, where application of micromolar amounts of CKs had a protective effect against *B. cinerea* and JA signaling was needed for this response (15). Thus, similarly to what happens with responses to biotrophic pathogens, increased CK content in the *s35* triple mutant amplifies JA-regulated defense responses to necrotrophic pathogens upon infection, suggesting that CK may be involved in this biotrophic-necrotrophic resistance tradeoff. Moreover, these results suggest that even though the effects of the *ckx3* and *ckx5* mutations are largely localized in reproductive tissue, these mutations can also affect necrotrophic and biotrophic defense responses in other parts of the plant such as leaves. Whether this outcome is also influenced by redox changes and ROXYs is unknown, but it is interesting to note that *ROXY*s are transcriptionally responsive to pathogen attack and SA and JA treatments (66, 67), and *roxy* mutants display altered susceptibility to *B. cinerea* (67, 68), and TGA activity is also susceptible to modification by overexpression of *ROXY* genes (67, 69). Thus, in addition to regulating SAM function, ROXYs may facilitate the crosstalk between SA and JA through CK and be responsible for the increased resistance phenotype to biotrophs and necrotrophs observed in the *s35* mutant.

Altogether, we propose a model (Supplemental Figure 9) where increased immunity suppresses plant growth, in part by the negative action of the defense hormones JA and, predominantly, SA on CK-signaling. While we can’t discount that crosstalk with additional hormones may be involved, the diminished growth suppression when CK levels are elevated indicates that CK-regulated physiological levels are at play. This growth suppression can act at the level of meristems, as seen by the reduced SAM and RAM sizes and altered phyllotactic patterns of these genotypes. This change in meristematic activity seems to be associated with an increased accumulation of ROS in reproductive tissues, which could be derived from changes in nitrogen signaling status for which CKs act as proxies, and possibly facilitated by the CC-type glutaredoxins (ROXYs), acting on yet unidentified target proteins. Increasing CK levels also changes the negative crosstalk of SA on JA signaling, generating plants that have increased resistance to both biotrophic and necrotrophic pathogens.

As it is progressively evident that breeding programs will need to adapt their practices for future sustainability, our research enables new avenues to be explored in crop species by exploiting phytohormones, natural signaling molecules governing plant fitness and survival, and their relationships through crosstalk. While other studies have uncovered instances of uncoupled growth and immunity (70–72), we hypothesized that the re-establishment of phytohormone homeostasis by phytohormone crosstalk engineering is the key to achieving this, bypassing costly and time-consuming efforts of identifying causal genes. The *s35* mutant in the model plant Arabidopsis we characterized here validates our hypothesis and allows for evaluation of the underlying mechanisms behind the CK-SA crosstalk that mediate synergistic growth and defense. Although this mutation combination yields similar fruiting capacity to wildtype, the reduced seed production is problematic. Yet, it is possible that this reduced seed set may be offset by fertilization that leads to larger seed size, as we have shown. If mimicked in crop species, this could still be an attractive phenotype due to a significant increase in resistance to a broad spectrum of pathogens, as well as the capacity to grow more individuals per area due to smaller plant size, potentially achieving a net harvest increase. Using the principles behind translational science, we envision that targeted phytohormone crosstalk modification can be applied to engineering improved crop species by precision genome editing of crosstalk players, to meet future crop yield demand under changing environmental conditions.

## MATERIALS AND METHODS

### Plant materials and growth conditions

All plant lines are in the Columbia-0 (Col-0) ecotype. *snc1-1* (10) and *ckx3-1 ckx5-1* (25) were obtained from ABRC. Plants were crossed, resulting in the *snc1 ckx3 ckx5* (*s35*) mutant, which was genotyped by PCR for homozygosity (see genotyping primers in Supplemental Table 2). Plants were grown in growth chambers in long-day conditions (16 h light/8 h dark, light intensity 120-150 µE, 22°C, RH 55% day/65% night) unless otherwise stated. For pathogen assays, plants were grown in short-day conditions (8 h light/16 h dark) with all other growth conditions kept consistent. For fertilization experiments, plants were given 1X Miracle-Gro Water Soluble All Purpose Plant Food (24-8-16) as needed.

### Phytohormone treatment

Phytohormone solutions were made using Sigma-Aldrich 6-benzylaminopurine (BA) and salicylic acid (SA). Dimethyl sulfoxide (DMSO) was used as the solvent. Spray treatments were done using Prevail sprayers. For liquid phytohormone treatment, seedlings were grown vertically on plates of 1x Murashige and Skoog (MS) with 1% sucrose and 0.6% agar for two weeks and transferred to liquid MS media for a 1 h acclimation period in growth chamber, shaking at 75 rpm. Phytohormone or DMSO control was then added to flasks to the final concentration indicated and seedlings harvested at specified times.

### Scanning electron microscopy

Primary shoot apical meristems of 6-week-old plants were hand dissected to stage 6 buds (27). Meristems were then fixed in 100% dry methanol for 10 minutes, then 100% dry ethanol for 30 minutes two times. Samples were critically dried with ethanol as the transitional fluid. Meristems were mounted upright on SEM stubs and gold coated. Images were captured using JEOL JSM-6500 Field Emission Scanning Electron Microscope at Colorado State University.

### RNA isolation and qRT-PCR

All plant tissue types were ground frozen using a tissue lyser and total RNA was extracted using the Qiagen RNeasy Mini Kit. RNA quality was assessed by A_260_/A_280_ and A_260_/A_230_ ratios via a NanoDrop. RNA samples of acceptable quality were treated with Invitrogen TURBO DNase as per the manufacturer’s instructions. DNase-treated RNA was checked for genomic DNA contamination by qRT-PCR using primers for AT5G66770. cDNA was synthesized using Quantabio QScript per the manufacturer’s instructions. cDNA was checked for full extension using three *PP2A* (AT1G13320) amplicons 1 kB apart. cDNA samples with C_q_ differences between primer pairs below 1.5 were acceptable for qRT-PCR gene expression. All qRT-PCR was performed using Quantabio PerfeCTa SYBR Green FastMix on Bio-Rad CFX Connect Real-Time PCR System, with CFX Maestro Software used for analysis. *UBIQUITIN 10* (AT4G05320) was used as a reference gene in all experiments. All qRT-PCR primer sequences are provided in Supplemental Table 3.

### RNA sequencing

All samples were collected developmentally, when the primary shoot had at least one fully expanded flower (stage 13+) present (27), following a modified protocol from (32). Inflorescences were dissected to stage 12 and under and cut from the stem. Between 15-18 inflorescences were pooled per sample. Samples were ground frozen using a tissue lyser and total RNA was extracted using the Qiagen RNeasy Micro Kit. RNA quality was assessed by A_260_/A_280_ and A_260_/A_230_ ratios via a NanoDrop and RIN_e_ scores via an Agilent 4150 TapeStation. RNA was sequenced using Illumina technology. Raw reads were filtered using fastp (73) by removing reads containing adapters, reads containing N > 10%, and low-quality reads. Paired-end clean reads were mapped to the TAIR10 reference genome (74) using HISAT2 software (75). RPKM of each gene was calculated using featureCounts (76). Benjamini and Hochberg’s approach for controlling the False Discovery Rate (FDR) by the DESeq2 R package (77) was used to determine genes with an adjusted P value < 0.05 and were assigned as differentially expressed. Gene ontology (GO) enrichment analysis of differentially expressed genes was performed using PANTHER 17.0 database [http://go.pantherdb.org/; (33)]. GO terms with an FDR-corrected P value < 0.05 were considered significantly enriched. GO graphs in Figure 5E through 5G were created using the modified R code from (78).

### *Hyaloperonospora arabidopsidis* inoculation and trypan blue staining

*Hyaloperonospora arabidopsidis* Noco2 (*Hpa*) was propagated on the susceptible Col-0 accession. Two-week-old plants were sprayed with *Hpa* spores (5×10^4^/mL) using a pressurized sprayer (Preval). Inoculated plants were covered with a transparent plastic dome to maintain high humidity. One day after the first appearance of sporangiophores (6 dpi) the first pair of true leaves was collected from three individual plants and added to a previously weighed 1.7 mL microcentrifuge tube containing 300 µL of sterile water, for a total of six leaves per sample, and weighed again to determine fresh weight. Spores were counted using a hemacytometer. Spore counts from at least four samples per genotype were determined.

For trypan blue staining, plants were harvested at 4 dpi and stained with a 3:1 ethanol:lacto-phenol trypan blue solution [1:1:1:1 phenol: lactic acid:water: glycerol and 0.05% trypan blue (Sigma-Aldrich)], at 95°C, for 5 minutes, and moved to room temperature for 10 minutes. Excess staining was removed with 15 M chloral hydrate (Sigma-Aldrich). Samples were moved to 50% glycerol for storage and mounting. Pictures were taken with a Nikon DS-Fi2 Microscope CCD camera mounted on a Nikon SMZ18 stereomicroscope.

### Hydrogen peroxide staining

Using a modified protocol from (79), rosettes and primary inflorescences from 6-week-old plants were stained with 1 mg/mL 3,3-diaminobenzidine (DAB) solution overnight on a laboratory shaker. The next day, samples were cleared with 3:1:1 enthanol: acetic acid: glycerol bleaching solution overnight on a laboratory shaker. The bleaching solution was replaced as needed. After clearing all chlorophyll, samples were preserved in 50% glycerol. Pictures were taken with a Dino-Lite Edge^Plus^ 1.3MP AM4917 Series microscope.

### TCSn::GFP imaging

Wildtype and transgenic *TCSn::GFP* seedlings (22) were grown vertically on MS plates supplemented with either water (mock) or 50 µM SA for 10-14 days. Seedling roots were mounted on slides and imaged using a Leica DM 5000-D fluorescence optical microscope.

### Pseudomonas syringae inoculation

Plants were germinated on pots covered with a plastic mesh. Two-week-old seedlings were watered in the morning of the inoculation day. Plants were dip-inoculated with a bacterial suspension as described by (80) with noted changes. *Pseudomonas syringae* pv. *tomato* DC3000 EV (*Pst*) was grown on King’s B (KB) Media supplemented with rifampicin (50 µg/mL) and kanamycin (50 µg/mL) and a bacteria suspension made in 10 mM MgCl_2_ for a concentration of 1×10^5^ CFU/mL was used for plant inoculation. Seedlings were collected and ground in 10 mM MgCl_2,_ and serial dilutions were used to determine the CFU/mg fresh weight (FW). Day 0 dilutions were plated on KB plates containing kanamycin and rifampicin, and day 3 dilutions were plated on KB plates containing rifampicin (50 µg/mL) and cycloheximide (50 µg/mL).

### Botrytis cinerea inoculation

*Botrytis cinerea* BO5.10 (*B. cinerea*) was grown on 0.5x Potato Dextrose Agar (PDA) plates until sporulation. A spore solution of 0.5×10^4^ spores/mL was prepared in ½ strength organic grape juice (R.W. Knudson Family Organic Juice, Just Concord) and 0.05% tween. The spore solution was sprayed on 6-week-old plants using a Prevail sprayer. Plants were placed in a flat under a water-sprayed plastic dome to maintain humidity. Pictures of plants were taken at 96 hpi. Plants were analyzed by the percentage of necrotic tissue of the rosette, and each plant was qualitatively assigned to one of the five disease index categories (20%, 40%, 60%, 80%, and 100% of leaves displaying disease symptoms). Disease indexes were averaged per genotype and a two-way ANOVA analysis with biological replicate as a blocking factor was performed to evaluate significance.

### Phytohormone quantification

For SA and JA quantification, leaf tissue from 6-week-old plants, or primary inflorescences from 8-week-old plants were dissected to stage 15 buds (27), were collected. All samples were immediately flash-frozen in liquid nitrogen after harvest and later lyophilized. Leaves from three plants (or approximately 30 inflorescences) were pooled per replicate. Tissue samples were homogenized and extracted with organic solvents, and LC-MS/MS analysis was performed on a Waters Acquity Classic UPLC coupled to a Waters Xevo TQ-S triple quadrupole mass spectrometer at the Colorado State University Bioanalysis and Omics (ARC-BIO) facility (81).

For CK quantification, primary inflorescences were collected as described above. Sample extraction and cytokinin quantification using the methods described in (82). A Waters Xevo TQ-S quantitative mass spectrometer and Waters Acquity UPLC HSS T3 (1.8 µm, 2.1 x 100 mm) column were used.

## Author contributions

G.A.J. and C.T.A. planned the research. H.M.B. made the crosses and conducted the *TCSn::GFP* experiments. H.S. and M.K. performed the CK quantification. Colorado State University Bioanalysis and Omics (ARC-BIO) facility performed the SA and JA quantification. G.A.J. conducted all remaining experiments and performed the data analyses. C.T.A. supervised the experiments. G.A.J. and C.T.A. wrote the manuscript with contributions from all authors.

## Supplemental figures

**Supplemental Figure 1:**
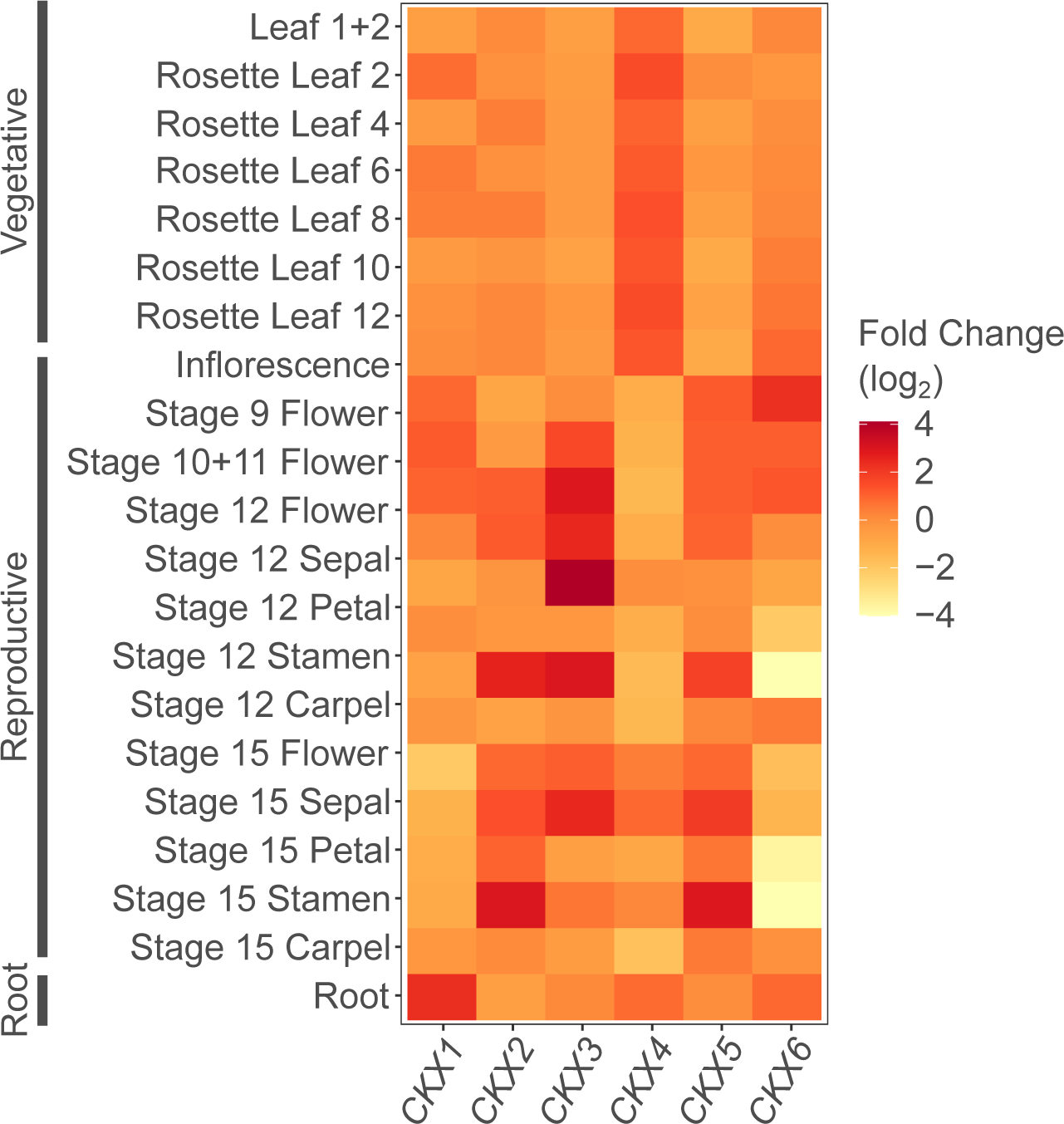
Tissue specificity of *CKX* gene expression. Publicly available gene expression data of six out of the seven *CKX* genes in different plant tissues. Expression values are log_2_-transformed. Data acquired and analyzed using the e-northern tool of the Bio-Analytic Resource for Arabidopsis Functional Genomics [http://bar.utoronto.ca; (83)]. Microarray gene expression normalized with robust multichip average (RMA) method.

**Supplemental Figure 2:**
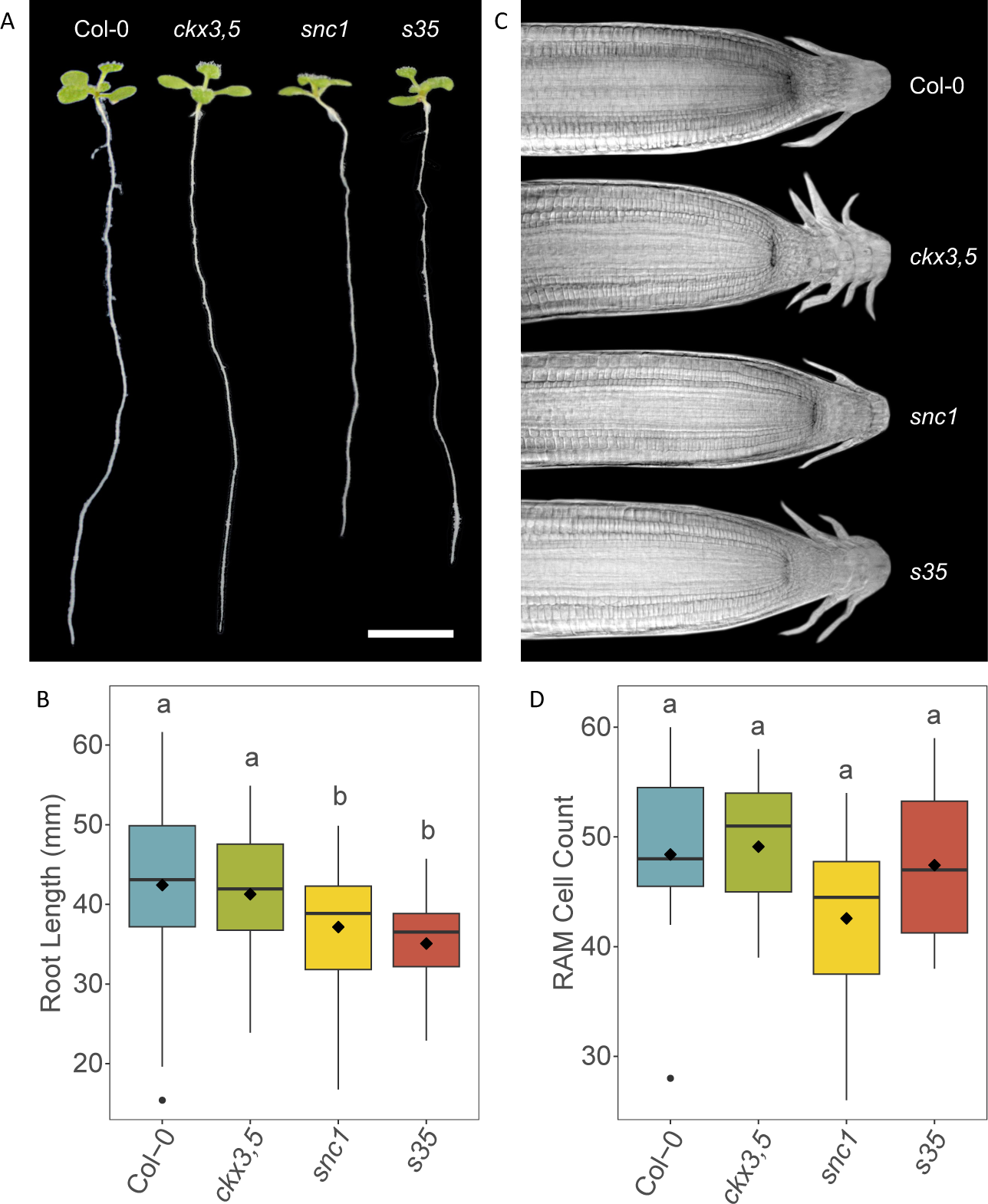
*s35* root phenotype is similar to that of *snc1*. (**A**) Representatives of roots grown on vertical MS plates. Scale bar = 1 cm. (**B**) Root length 9 days post-germination. Data pooled from three independent biological replicates are shown. n ≥ 48. Letters indicate differences at a P < 0.05 significance level using a one-way ANOVA analysis with biological replicate as a blocking factor and Tukey post-hoc correction. (**C**) Representatives of root tips. (**D**) The number of cortex cells in the zone of cell proliferation. n ≥ 10. Letters indicate differences at a P < 0.05 significance level using a one-way ANOVA analysis with biological replicate as a blocking factor and Tukey post-hoc correction. For (**D**), letters indicate differences at a P < 0.05 significance level using a one-way ANOVA analysis with a Tukey post-hoc correction.

**Supplemental Figure 3:**
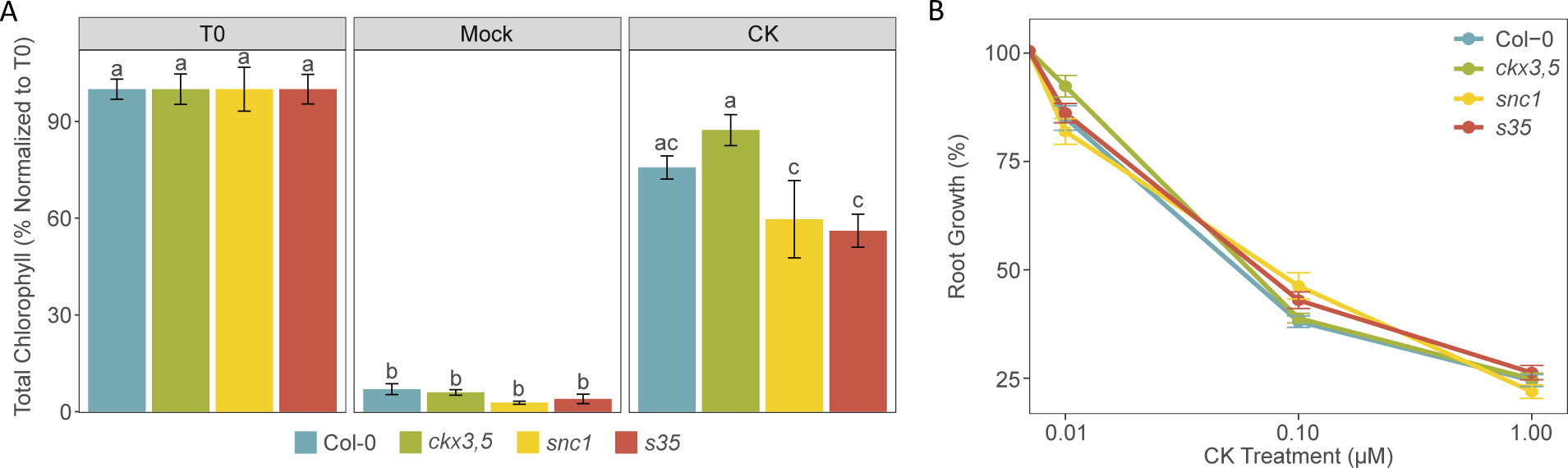
*s35* sensitivity to CK is not altered in vegetative tissues. (**A**) Total chlorophyll (*a* + *b*) content of mature leaves normalized to non-treated samples (T0) after mock or CK treatment. Leaves from 6-week-old plants were at the petiole and placed in the dark in either DMSO mock or 100 nM CK solutions for 7 days in the dark to allow for senescence. Data pooled from two independent biological replicates are shown. (**B**) Percentage of root length after growth on three concentrations of CK normalized to DMSO mock. Sterilized seeds were germinated on vertical MS plates with either DMSO, 0.01 µM CK, 0.1 µM CK, or 1 µM CK. Roots were measured 10 days post germination. n > 34. Data pooled from three independent biological replicates are shown. For (**A**) and (**B**), error bars represent s.e.m. and letters indicate differences at a P < 0.05 significance level using a two-way ANOVA analysis with a Tukey post-hoc correction. For (**B**), the differences between the slopes of each genotype were not significant.

**Supplemental Figure 4:**
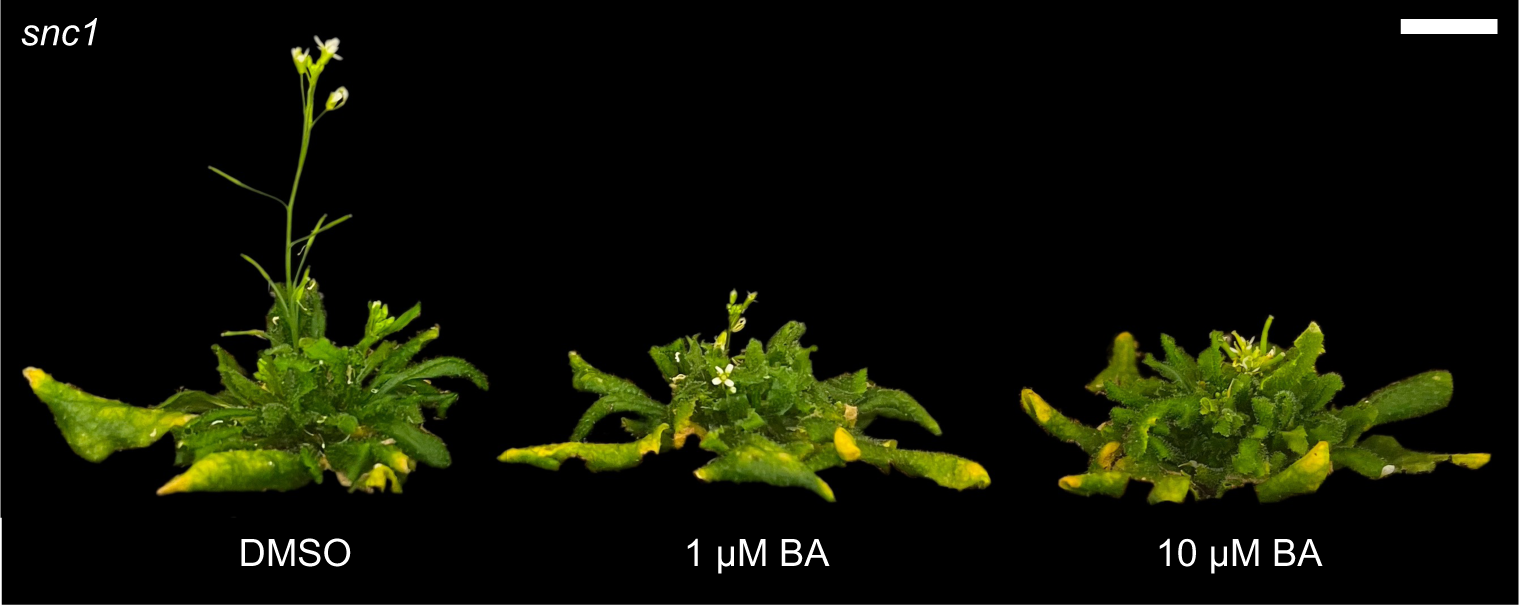
Dose-dependent treatment with cytokinin of *snc1* plants partially phenocopies the *s35* inflorescence morphology. *snc1* was sprayed with either 0.1% dimethyl sulfoxide (DMSO; left), 1 µM 6-benzylaminopurine (BA; middle), or 10 µM BA (left) in water. Plants were spray-treated once per week, starting at 3 weeks old for a total of 5 treatments. Representatives of 10-week-old plants shown. At least two independent biological replicates of the experiment were conducted with similar results. n = 9. Scale bar = 1 cm.

**Supplemental Figure 5:**
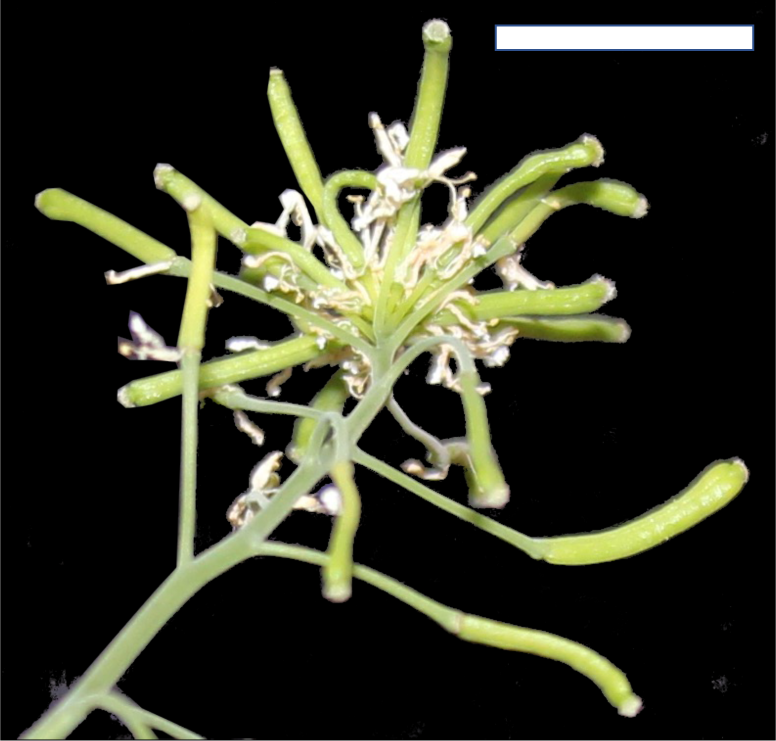
*cpr1 ckx3 ckx5* (*c35*) has a starburst inflorescence phenotype similar to that of *s35*. The *c35* cross was confirmed via genotyping for homozygosity. Representative shown. Scale bar = 1 cm.

**Supplemental Figure 6:**
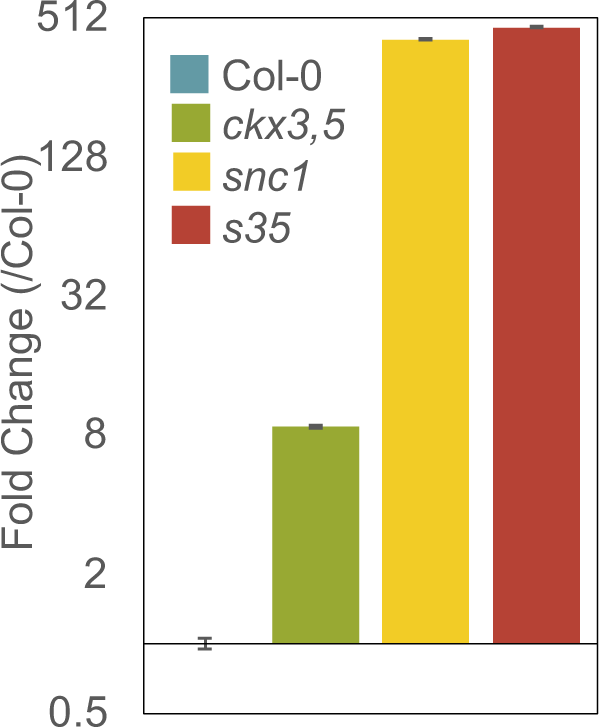
The *s35* triple mutant displays constitutive expression of *PR1*, similar to *snc1*. Basal levels of *PATHOGENESIS-RELATED 1* (*PR1*) transcripts were determined by qRT-PCR relative to wildtype plants and normalized to *UBIQUITIN 10* (*UBQ10*). Error bars represent s.e.m. from three biological replicates and correspond to upper and lower limits of 95% confidence intervals. For better visualization, the axis is log_2_-transformed. Data pooled from three independent biological replicates are shown.

**Supplemental Figure 7:**
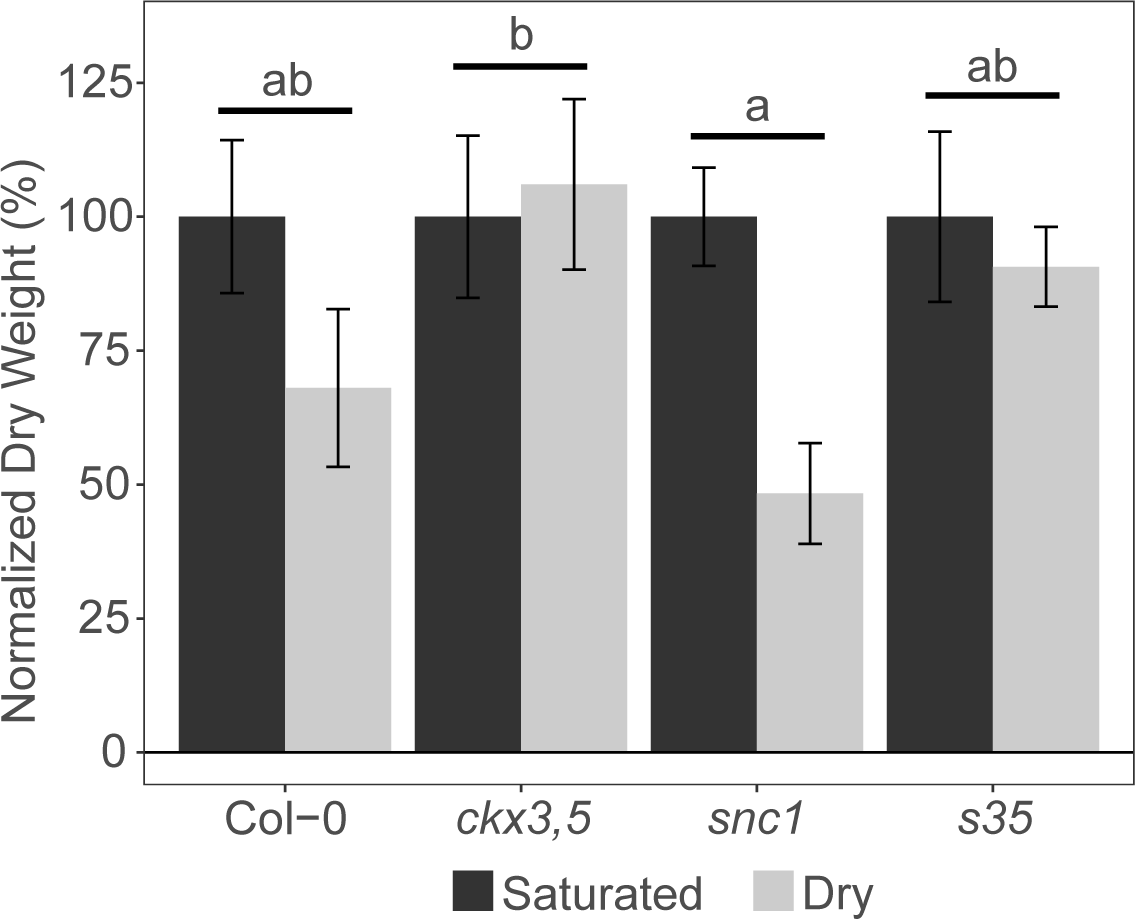
The *s35* triple mutant has comparable drought tolerance as wildtype. Plants were grown as per the soil moisture deficit treatment protocol described in (84). Plants were kept well-watered at 100% water capacity (Saturated) or underwent dry-down to 40% water capacity for 7 consecutive days (Dry). Rosette tissue was harvested, dried, and weighed. Tissue weights were normalized to the saturated control treatment within genotype. Error bars represent s.e.m. and correspond to upper and lower limits of 95% confidence intervals. Data from one out of two total independent biological replicates shown. Letters indicate differences between the treatments at a P < 0.05 significance level using a two-way ANOVA analysis.

**Supplemental Figure 8:**
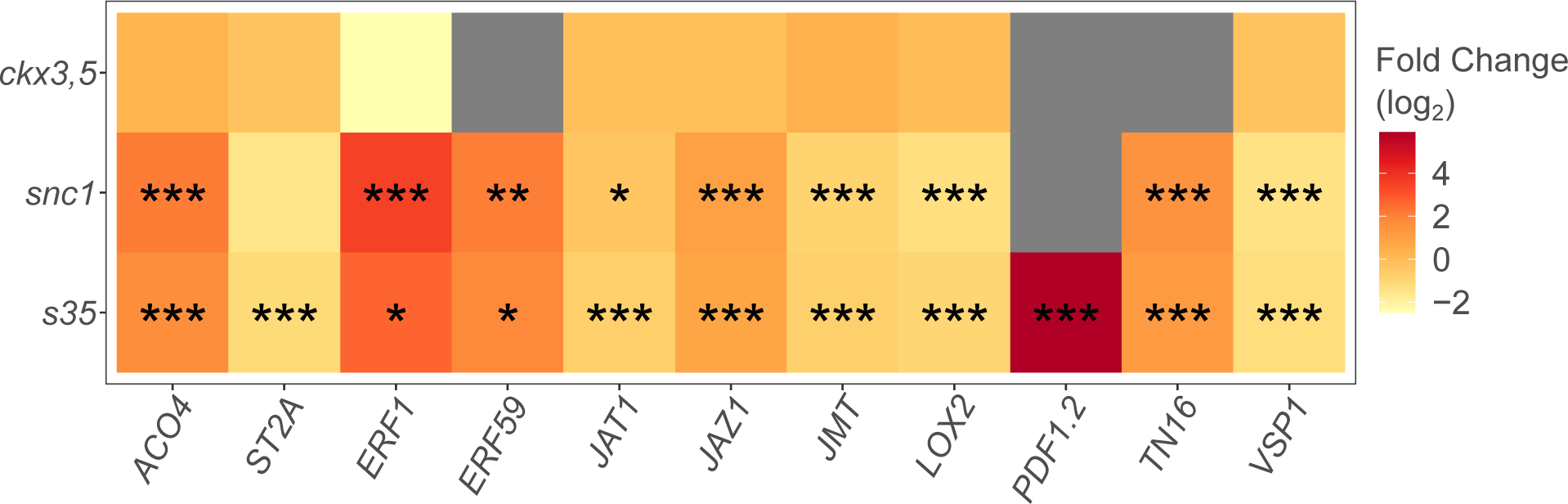
*s35* and *snc1* inflorescences display marginally altered expression of JA-regulated genes. Heatmap of most differentially expressed JA-regulated genes in the *s35* mutant, and its parental genotypes, in relation to wildtype Col-0 plants. The gene list was assembled from relevant literature and searched against the significantly DEG of each genotype from the transcriptomic analysis. Statistically significant differences from wildtype inflorescences (FDR) are represented by asterisks (* = P value < 0.05, ** = P value < 0.01, *** = P value < 0.001). Gray color means absence of differential expression in dataset.

**Supplemental Figure 9:**
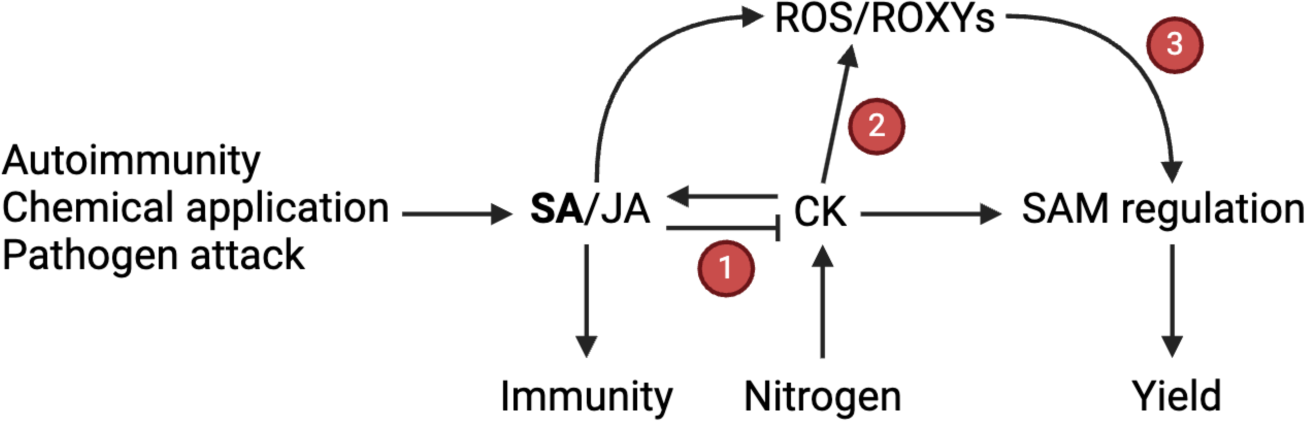
Proposed model of SAM regulation in relation to the CK-SA crosstalk. Upon immunity activation, JA, and mostly SA, repress CK signaling **(1)**. Immunity activation also triggers the production of reactive oxygen species (ROS), which are also under control of CK (**2**). Altered redox status modulates SAM activity, possibly through the action of CC-type glutaredoxins (ROXYs) oxidoreductive modification of target proteins (**3**). CKs act as a long-range signal for nitrate availability, affecting SAM activity, and promoting reproductive growth, which may also happen by the action of the ROXY proteins. Numbers represent relationships supported or suggested by our data. Model created with BioRender.com.

**Supplemental Table 1:**
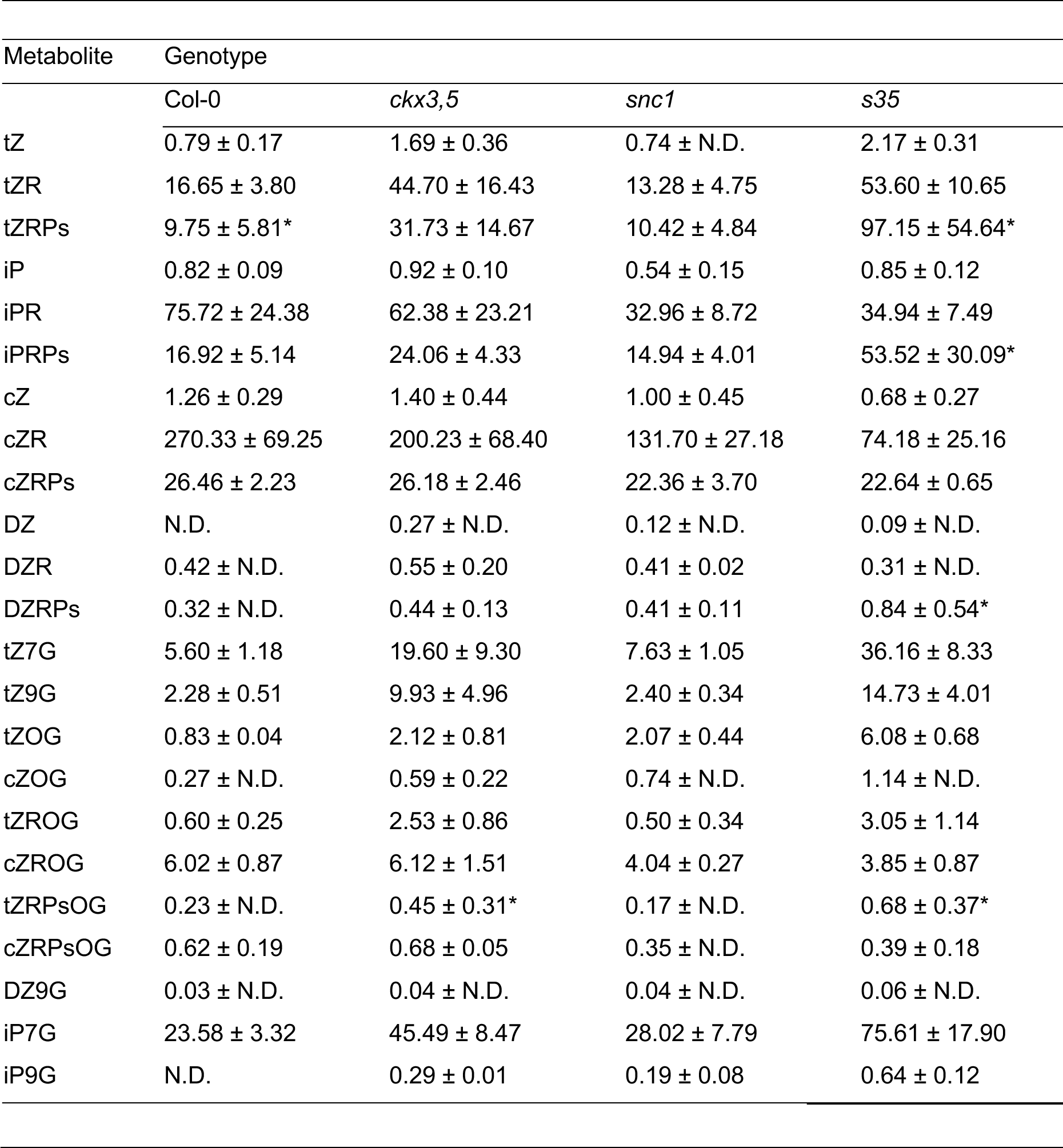
The *s35* mutant has a unique CK species profile. Approximately 30 inflorescences per sample were harvested and pooled from 8-week-old plants. All samples were lyophilized. Three independent biological samples were harvested from each genotype. Data shown are pmol/g dry weight ± s.d.; n = 3. Asterisks (*) indicates percent s.d. over 50; peak annotations were rechecked. N.D. = not detected.

| Metabolite | Genotype |  |  |  |
| --- | --- | --- | --- | --- |
|  | Col-0 | *ckx3,5* | *snc1* | *s35* |
| tZ | 0.79 ± 0.17 | 1.69 ± 0.36 | 0.74 ± N.D. | 2.17 ± 0.31 |
| tZR | 16.65 ± 3.80 | 44.70 ± 16.43 | 13.28 ± 4.75 | 53.60 ± 10.65 |
| tZRPs | 9.75 ± 5.81* | 31.73 ± 14.67 | 10.42 ± 4.84 | 97.15 ± 54.64* |
| iP | 0.82 ± 0.09 | 0.92 ± 0.10 | 0.54 ± 0.15 | 0.85 ± 0.12 |
| iPR | 75.72 ± 24.38 | 62.38 ± 23.21 | 32.96 ± 8.72 | 34.94 ± 7.49 |
| iPRPs | 16.92 ± 5.14 | 24.06 ± 4.33 | 14.94 ± 4.01 | 53.52 ± 30.09* |
| cZ | 1.26 ± 0.29 | 1.40 ± 0.44 | 1.00 ± 0.45 | 0.68 ± 0.27 |
| cZR | 270.33 ± 69.25 | 200.23 ± 68.40 | 131.70 ± 27.18 | 74.18 ± 25.16 |
| cZRPs | 26.46 ± 2.23 | 26.18 ± 2.46 | 22.36 ± 3.70 | 22.64 ± 0.65 |
| DZ | N.D. | 0.27 ± N.D. | 0.12 ± N.D. | 0.09 ± N.D. |
| DZR | 0.42 ± N.D. | 0.55 ± 0.20 | 0.41 ± 0.02 | 0.31 ± N.D. |
| DZRPs | 0.32 ± N.D. | 0.44 ± 0.13 | 0.41 ± 0.11 | 0.84 ± 0.54* |
| tZ7G | 5.60 ± 1.18 | 19.60 ± 9.30 | 7.63 ± 1.05 | 36.16 ± 8.33 |
| tZ9G | 2.28 ± 0.51 | 9.93 ± 4.96 | 2.40 ± 0.34 | 14.73 ± 4.01 |
| tZOG | 0.83 ± 0.04 | 2.12 ± 0.81 | 2.07 ± 0.44 | 6.08 ± 0.68 |
| cZOG | 0.27 ± N.D. | 0.59 ± 0.22 | 0.74 ± N.D. | 1.14 ± N.D. |
| tZROG | 0.60 ± 0.25 | 2.53 ± 0.86 | 0.50 ± 0.34 | 3.05 ± 1.14 |
| cZROG | 6.02 ± 0.87 | 6.12 ± 1.51 | 4.04 ± 0.27 | 3.85 ± 0.87 |
| tZRPsOG | 0.23 ± N.D. | 0.45 ± 0.31* | 0.17 ± N.D. | 0.68 ± 0.37* |
| cZRPsOG | 0.62 ± 0.19 | 0.68 ± 0.05 | 0.35 ± N.D. | 0.39 ± 0.18 |
| DZ9G | 0.03 ± N.D. | 0.04 ± N.D. | 0.04 ± N.D. | 0.06 ± N.D. |
| iP7G | 23.58 ± 3.32 | 45.49 ± 8.47 | 28.02 ± 7.79 | 75.61 ± 17.90 |
| iP9G | N.D. | 0.29 ± 0.01 | 0.19 ± 0.08 | 0.64 ± 0.12 |

**Supplemental Table 2:**
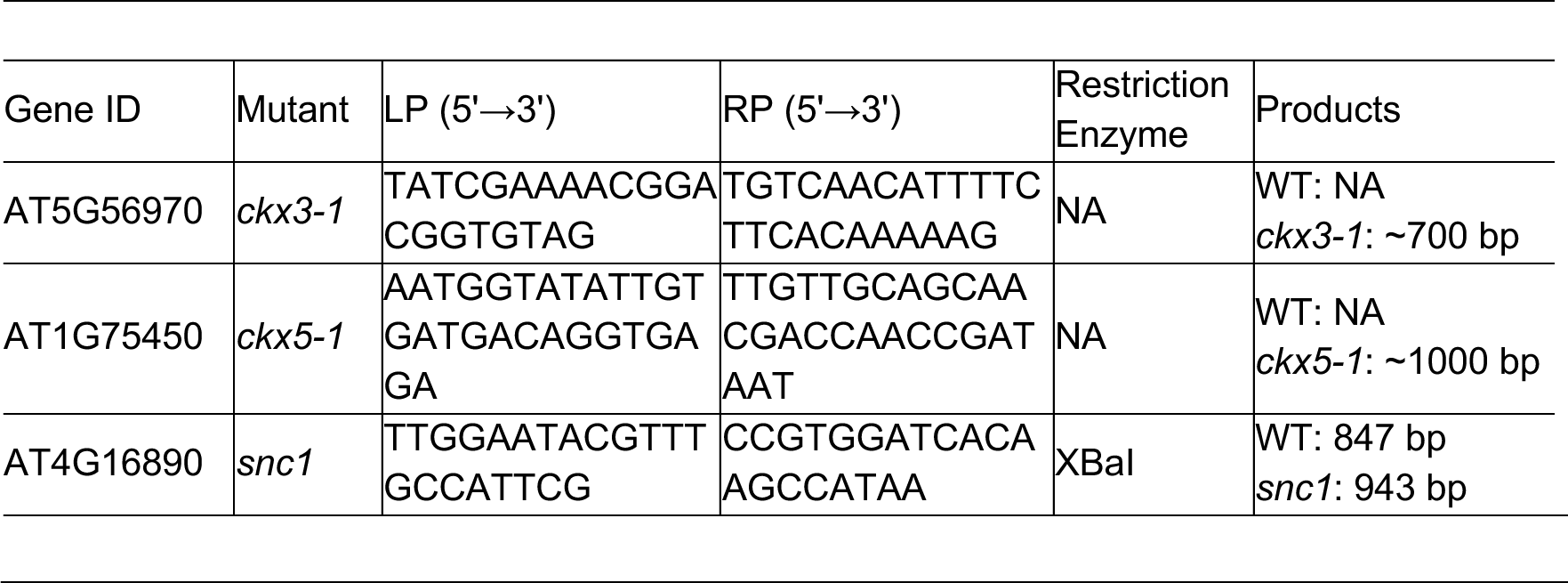
Primers and restriction enzymes for mutant genotyping.

**Supplemental Table 3:**
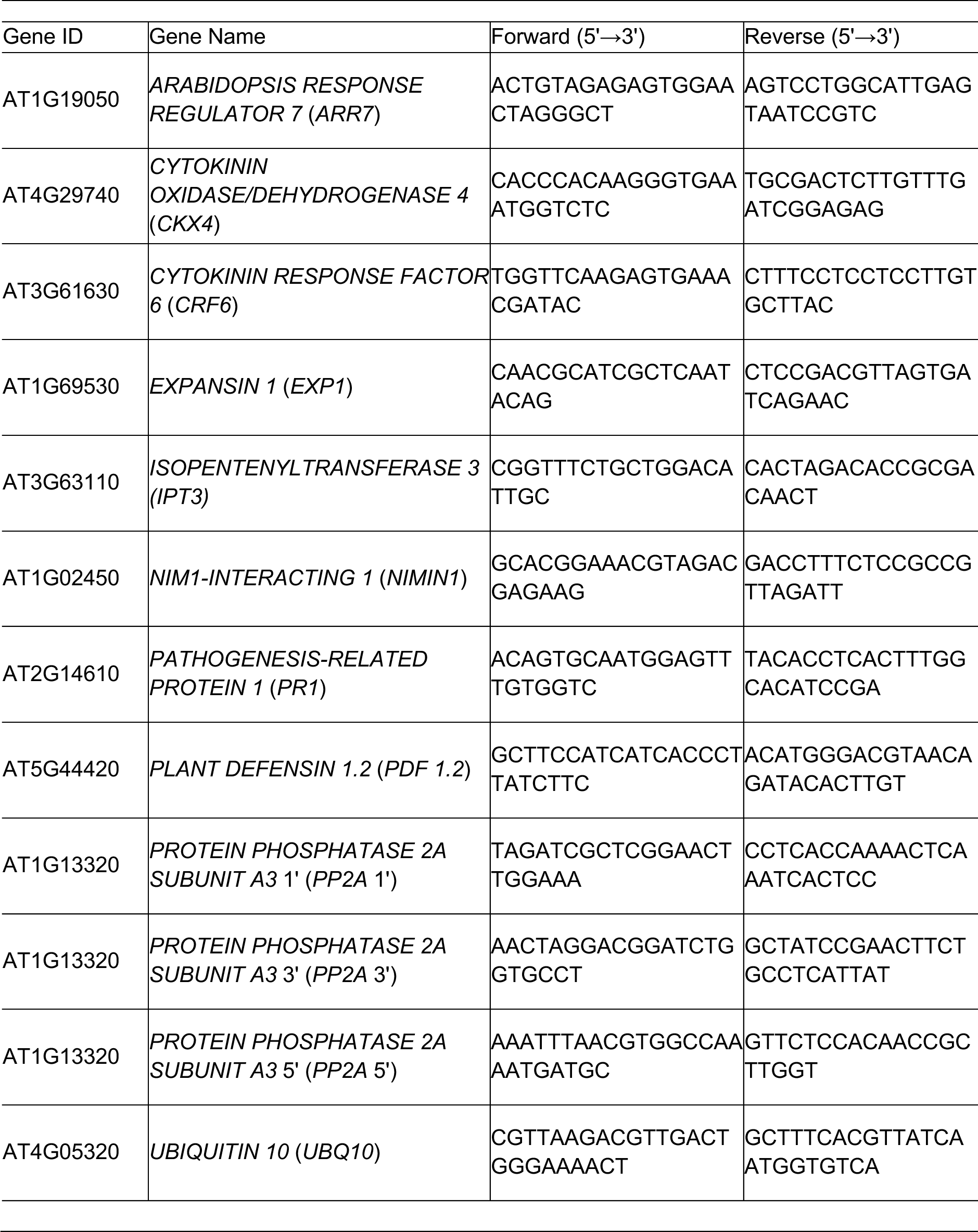
qRT-PCR quality control and gene-specific primers for gene expression analysis.

